# A toolbox of FRET-based c-di-GMP biosensors and its FRET-To-Sort application for genome-wide mapping of the second messenger regulatory network

**DOI:** 10.1101/2024.08.21.609041

**Authors:** Liyun Wang, Gabriele Malengo, Ananda Sanches-Medeiros, Xuanlin Chen, Nataliya Teteneva, Silvia González Sierra, Ming C. Hammond, Victor Sourjik

**Affiliations:** Max Planck Institute for Terrestrial Microbiology & Center for Synthetic Microbiology (SYNMIKRO), Karl-von-Frisch-Straße 14, 35043 Marburg, Germany; Department of Chemistry and Henry Eyring Center for Cell & Genome Science, University of Utah, Salt Lake City, Utah 84112, United States; Department of Chemistry, University of California, Berkeley, California 94720, United States

## Abstract

C-di-GMP is a widespread second messenger, coordinating various cellular functions in bacteria. Levels of c-di-GMP can be highly dynamic and vary over a wide range of concentrations. Here we constructed a large set of FRET-based c-di-GMP biosensors, using homologues of c-di-GMP-binding effector YcgR from different bacterial species. This biosensor library was characterized using a newly established protocol to quantify FRET efficiency using flow cytometry. The resulting toolbox of 18 selected biosensors that undergo large FRET signal change upon c-di-GMP binding displays a ∼100-fold range of c-di-GMP binding affinities. We combined this toolbox with a barcoded Tn5 transposon library and cytometry-based cell sorting to develop FRET-To-Sort, a new application for systematic characterization of gene networks regulating levels of FRET-detected small molecules. Applied to planktonic *E. coli* cells, FRET-To-Sort identified both known and novel regulatory modules controlling c-di-GMP levels, including flagellum and fimbria biogenesis, lipid metabolism and stress response genes.

---

The second messenger cyclic diguanosine monophosphate (c-di-GMP), now recognized as a ubiquitous signaling molecule in bacteria, regulates a multitude of cellular processes that are primarily related to lifestyle transitions, including switch from motility to sessility, virulence, biofilm formation and cell cycle control ^1,2^. In most bacterial species, c-di-GMP levels are controlled by a highly complex network of multiple diguanylate cyclases (DGCs) and phosphodiesterases (PDEs), which are responsible for the synthesis and the hydrolysis of c-di-GMP, respectively. These enzymes can be responsive to environmental and cellular cues, mediating signaling via c-di-GMP binding effectors, such as transcription factors ^3,4^, riboswitches ^5^ or post-translational regulators ^6–8^. A large class of the latter are proteins that contain PilZ domains, which are activated by c-di-GMP and generally function via protein-protein interactions, exemplified by the flagellar brake protein YcgR in *E. coli* ^7,8^, a stand-alone PilZ domain.

Monitoring c-di-GMP dynamics is thus critical to understand the bacterial cell physiology. For that purpose, several c-di-GMP biosensors have been developed and employed over the last decade, such as transcriptional biosensors, in which a c-di-GMP-responsive promoter controls expression of the green fluorescent protein (GFP) gene ^9–11^. However, while being powerful in visualizing changes of c-di-GMP over generations, transcriptional reporters do not allow monitoring real-time dynamics, and their activity could be potentially sensitive to other cellular factors even when decoupled from regulation by c-di-GMP ^12^. Hence, several different direct biosensors of c-di-GMP were constructed, including RNA-based biosensors where a c-di-GMP-binding riboswitch aptamer was fused to the fluorescent-dye binding Spinach aptamer ^13,14^, bimolecular complementation for a split fluorescent protein fused to dimerizing c-di-GMP-binding protein BldD ^15^, and a few biosensors based on Förster resonance energy transfer (FRET) that utilized c-di-GMP PilZ-domain proteins fused to FRET pairs composed by donor and acceptor fluorescent proteins ^16,17^.

While the above-mentioned individual biosensors were highly useful to determine c-di-GMP levels *in vivo*, all of them are limited to a relatively narrow range, typically one decade, of c-di-GMP concentrations. In contrast, intracellular c-di-GMP can vary strongly among bacterial species and strains ^18,19^, between sessile and motile lifestyles ^2,20,21^, during cell cycle ^16,20,22^ as well as upon changes of environmental factors ^23^, such as nutrients ^24,25^, oxygen ^25^ and antibiotics ^26^, spanning the range from <50 nM to a few μM ^1^. Even in the same organism under same culture condition, levels of c-di-GMP can vary over a wide range dependent on the growth phase, as well as among cells in the same population ^1,23,25,27^.

Thus, to enable investigation of intracellular c-di-GMP levels under different conditions and/or in different species and strains, we constructed a set of biosensors with different affinities using PilZ-domain proteins. We adapted a library of 90 PilZ-domain containing proteins, homologous to *E. coli* YcgR, that was previously established for the *in vitro* measurement of c-di-GMP levels using bioluminescence resonance energy transfer (BRET) ^28^, and converted it into *in vivo* FRET biosensors. The response magnitude and c-di-GMP binding affinity of individual sensors were then characterized using a newly established high-throughput FRET analysis using flow cytometry. This resulted in a toolbox of c-di-GMP FRET biosensors exhibiting large signal changes and different affinities, that could be used for a variety of applications. We subsequently harnessed sensors from our toolbox to establish a new <u>FRET</u>-based screening approach for genome-wide identification of mutants with high or low c-di-GMP levels in a barcoded Tn5 transp<u>o</u>son library using fluorescence-activated cell <u>sort</u>ing (FRET-To-Sort). Its application to *E. coli* provided insights into the complex c-di-GMP regulatory network, which prominently included flagellum and fimbria biogenesis, lipid metabolism and stress response.

## Results

### Construction and characterization of the FRET-based c-di-GMP biosensor library

Upon c-di-GMP binding, PilZ domains can undergo a conformational change leading to an altered distance between their termini ^6,29^. This enables their use as FRET-biosensors by fusing their termini to donor and acceptor fluorophores. Because the efficiency of energy transfer from the donor to the acceptor is inversely proportional to the sixth power of the distance between these two fluorophores and this transfer occurs within nanoseconds ^30^, such FRET biosensors can exhibit highly sensitive and rapid response to the concentration change of c-di-GMP. Nevertheless, the extent of the conformational change and affinity of c-di-GMP binding may vary between different PilZ domain proteins ^28,31^. In order to construct a set of FRET-based biosensors with different affinities and large conformational change, we therefore used the previously assembled library of 90 *ycgR* sequences from different organisms ^28^, fusing each YcgR homologue to mNeonGreen (mNG, FRET acceptor) and mTurquoise2 (mTq2, FRET donor) fluorescent proteins (Fig. 1a and Supplementary Table 1). These biosensors were named according to their positions in the library, and tested for their conformational change (reflected by FRET efficiency change) upon c-di-GMP binding *in vivo*, using strains respectively deleted for genes encoding major DGC (*ΔdgcE*) or major PDE (*ΔpdeH*) that control the global pool of c-di-GMP in *E. coli* ^2^.

**Fig. 1.**
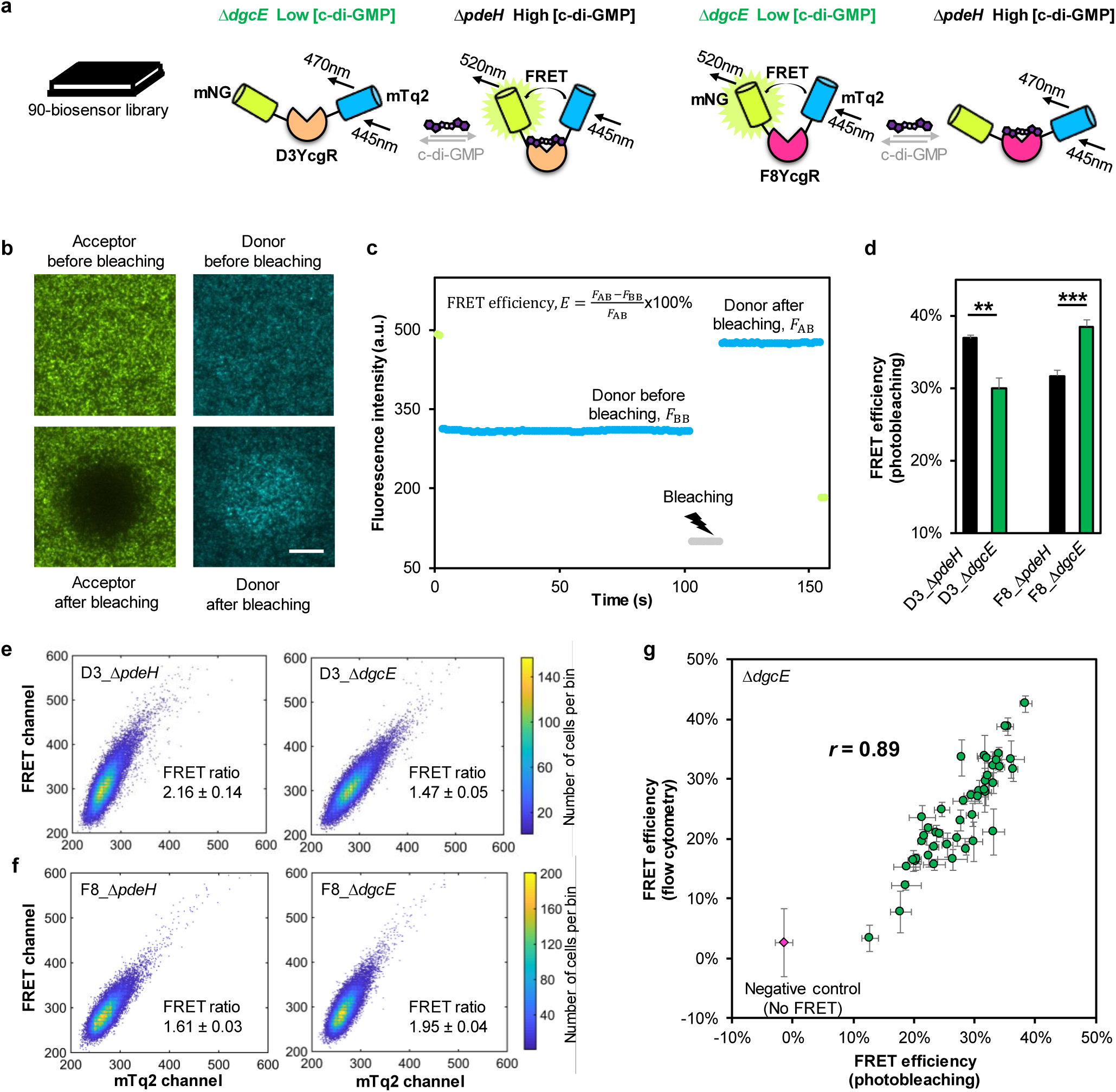
Construction and characterization of the c-di-GMP FRET biosensor library. **a,** Schematics of the FRET-based c-di-GMP biosensor library. To screen potential sensors for their sensitivity to changes in c-di-GMP, fusion proteins were transformed into *ΔpdeH* and *ΔdgcE* strains, which respectively have high or low levels of c-di-GMP, as indicated. Upon binding c-di-GMP, conformational change in biosensors might either decrease or increase the distance between the donor, mTurquoise2 (mTq2), and acceptor, mNeonGreen (mNG), fluorophores, resulting in respectively increased or decreased energy transfer from mTq2 to mNG. **b,c,** Quantification of FRET efficiency and effects of c-di-GMP using acceptor photobleaching microscopy, with *ΔpdeH* cells that express G2 biosensor shown as an example. Fluorescence of bacterial monolayer was imaged in donor and acceptor fluorescence channels, before and after acceptor photobleaching. Scale bar: 20 μm. FRET efficiency was calculated from acquired donor fluorescence as illustrated in (**c**). *F_AB_*, donor fluorescence after acceptor photobleaching. *F_BB_*, donor fluorescence before acceptor photobleaching. **d,** Values of FRET efficiency for D3 and F8 biosensors measured by acceptor photobleaching microscopy in *ΔpdeH* and *ΔdgcE* strains. \*\**P* <0.01; \*\*\**P* <0.001. *P* values were calculated using unpaired two-tailed *t-test*, n=3. Data are presented as mean ± SD. **e,f,** Measurements of sensitized-FRET using flow cytometry. Scatter plots show fluorescence in the donor (mTq2) and acceptor (FRET) channels of *ΔpdeH* or *ΔdgcE* cells expressing D3 (**e**) or F8 (**f**) biosensors, excited by 445 nm laser. Each dot presents one individual cell, and is colored by the number of cells per heat-map bin. The mean FRET ratio (the ratio of fluorescence intensity in the FRET channel over that in the donor channel) ± SEM is shown. n= 30000. **g,** Comparison of the values of FRET efficiency calculated using calibrated flow cytometry with those measured using acceptor photobleaching microscopy for different potential biosensors in *ΔdgcE* strain. The negative control corresponds to mTq2 and mNG co-expressed from different plasmids in *ΔdgcE* strain. n≥3. Data are presented as mean ± SD. Pearson correlation coefficient *r* is shown.

To validate the functionality of our biosensors, we first determined FRET efficiency by acceptor photobleaching ^30^ (Fig. 1b,c). Since fluorescence of the donor is quenched due to energy transfer to the acceptor, its unquenching upon photobleaching of the acceptor provides a direct measure of FRET efficiency. We observed that for several sensors, thus determined FRET efficiency showed significant differences between *ΔpdeH* (high c-di-GMP) or *ΔdgcE* (low c-di-GMP) background (Fig. 1d), supporting our expectation that they could be used as c-di-GMP biosensors. Notably, while some constructs (e.g., D3) showed higher FRET in *ΔpdeH* compared to *ΔdgcE* strain, others (e.g., F8) showed an opposite difference (Fig. 1d), suggesting that different homologues of YcgR can undergo opposite conformational changes after binding c-di-GMP (Fig. 1a).

While acceptor photobleaching microscopy allows straightforward measurements of FRET efficiency, this method is relatively laborious and not ideal for the high-throughput characterization of a large biosensor library in multiple strains/conditions. Therefore, we sought to establish a higher-throughput FRET assay using flow cytometry, based on the sensitized acceptor emission ^32^ (Extended Data Fig. 1a). Flow cytometry for *E. coli* cell cultures expressing D3 or F8 biosensors confirmed that upon donor excitation, the ratio of signals in the FRET channel (acceptor emission) and the donor emission channel (defined as FRET ratio) differed between *ΔpdeH* and *ΔdgcE* strains. FRET ratio increased at high level of c-di-GMP for D3 biosensor (Fig. 1e) and decreased at high level of c-di-GMP for F8 biosensor (Fig. 1f), thus consistent with the trend measured by acceptor photobleaching (Fig. 1d).

Nevertheless, although being a useful proxy, the FRET ratio calculated simply as the ratio between the sensitized-FRET and donor emission signals may respond nonlinearly to the change in FRET efficiency, and is instrument-and expression-dependent ^32^. We thus established a calibration for the flow cytometry data following previous method ^33^ to obtain a quantitative and accurate measure of FRET (Methods and Extended Data Fig. 1b). Validating this approach, FRET efficiency values for 46 randomly chosen biosensors expressed in *ΔdgcE* strain showed an excellent quantitative agreement between flow cytometry and photobleaching microscopy measurements (Fig. 1g), except a seeming offset between cytometry and microscopy data at low FRET efficiency (<10%).

### A toolbox of characterized c-di-GMP biosensors

We proceeded to characterize the entire library for FRET efficiencies in *ΔpdeH* (high c-di-GMP) and *ΔdgcE* (low c-di-GMP) strains using flow cytometry. Most of the biosensors displayed high FRET efficiency, with median values of ∼27% (Supplementary Fig. 1), which is consistent with proximity of the donor and acceptor fluorophores. Several biosensors showing low values of FRET efficiency (<10%) or high variability between replicates in both strains, were excluded from further analysis (Supplementary Fig. 2). For each of the remaining biosensors, we determined the ratio of FRET efficiencies in *ΔpdeH* and in *ΔdgcE* strains, which reflects the magnitude of their conformational change upon c-di-GMP binding (Fig. 2a). Most sensors showed a detectible difference (Fig. 2a), with a similar number of sensors showing either increased or decreased FRET at higher levels of c-di-GMP, corresponding to two possible conformational changes upon c-di-GMP binding (Fig. 1a). During this analysis, we noticed that the difference between FRET in *ΔpdeH* and *ΔdgcE* strains was generally smaller in the presence of 67 mM NaCl in the measurement buffer (Fig. 2a), which is discussed later.

**Fig. 2.**
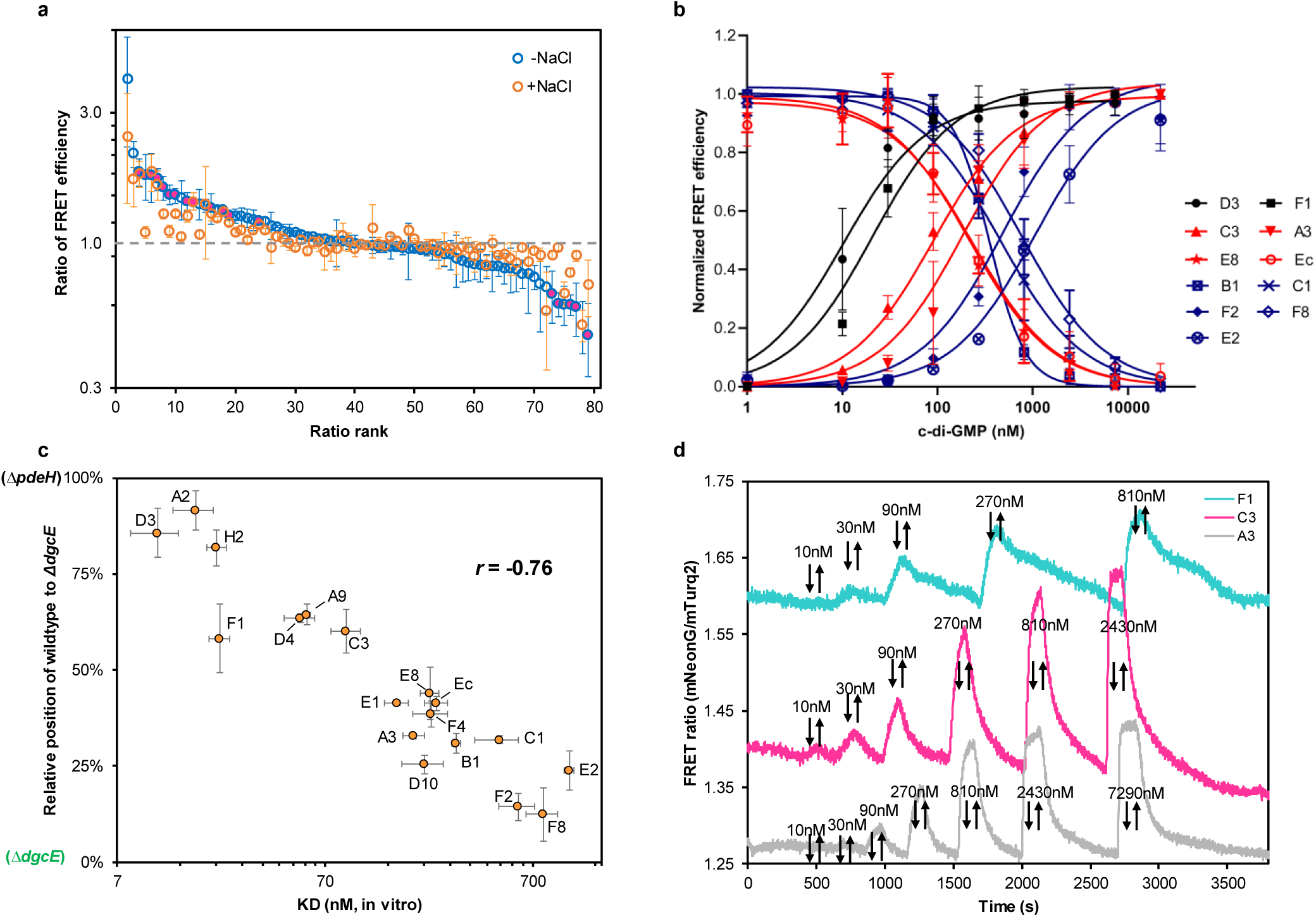
Establishment and characterization of the biosensor toolbox. **a,** Ranked ratio of the FRET efficiency for each biosensor in *ΔpdeH vs ΔdgcE* strains. Measurements were done either in presence of 67 mM NaCl (+NaCl) in the sampling buffer or in its absence (-NaCl), as indicated. Dashed line indicates no difference in FRET between the two strains. Magenta-filled data points indicate biosensors chosen for the toolbox. n=3. Data are presented as mean ± SD. **b,** Dose responses of selected biosensors chosen for the toolbox, measured in permeabilized *ΔdgcE* cells (i.e., *in vitro*). n≥3. Data are presented as mean ± SD. Black, red or blue color, respectively, indicates high, medium or low affinity. The curves for the other 7 biosensors in the toolbox are shown in Extended Data Fig. 2. **c,** *K*_D_ values, determined from *in vitro* dose-response measurements as in **b** and Extended Data Fig. 2, plotted against the relative FRET efficiency in wildtype relative to those in *ΔpdeH* and in *ΔdgcE* strains, which serves as a proxy of the *in vivo* sensor affinity (see Supplementary Fig. 3a, b). The *in-vivo* values were measured in the absence of NaCl. n≥3. Data are presented as mean ± SEM. Pearson correlation coefficient *r* is shown. **d,** Ratiometric FRET measurement of response to stepwise addition and subsequent removal of indicated concentrations of c-di-GMP, as indicated by arrows, in a flow chamber for selected biosensors in permeabilized *ΔdgcE* cells. FRET ratio was defined here as the ratio between fluorescence emission in the acceptor and in the donor channel while exciting donor fluorescence.

For further characterization, we chose 18 biosensors (Table 1) exhibiting top-ranked ratios of FRET efficiency between *ΔpdeH* and *ΔdgcE* strains and with the absolute levels of FRET efficiency above 15% in both strains. In order to determine the affinity of these selected biosensors to c-di-GMP, we rendered sensor-expressing *ΔdgcE* cells permeable to extracellular c-di-GMP by treating them with toluene ^34^. All 18 biosensors exhibited reproducible dose-dependent changes in FRET efficiency measured by flow cytometry upon incubation of such permeabilized *ΔdgcE* cells with extracellular c-di-GMP (Fig. 2b, Extended Data Fig. 2). The sign of their response – either increase or decrease in FRET upon c-di-GMP binding – was consistent with the *in vivo* difference between *ΔpdeH* and *ΔdgcE* strains (Table 1). The biosensors based on *E. coli* (Ec) and *Salmonella typhimurium* (C1) YcgR both displayed decreased FRET after binding c-di-GMP, which is consistent with literature ^16,35^. We observed an approximately 100-fold variation in sensor affinities to c-di-GMP, with the dissociation constant *K*_D_ ranging from ∼10 nM to ∼1 µM (Table 1). According to their *K*_D_ values, biosensors were classified in three groups of affinity, with *E. coli* YcgR being used as a reference for the definition of medium affinity. For a few YcgR homologues where *K*_D_ values were previously measured *in vitro* ^16,28,35,36^, the values of *K*_D_ determined here were generally similar, with moderate (up to 2.5-fold) discrepancies being likely due to the differences in methods, buffer compositions or the effect of fluorescent protein fusions on the binding affinity (Table 1).

**Table 1.**
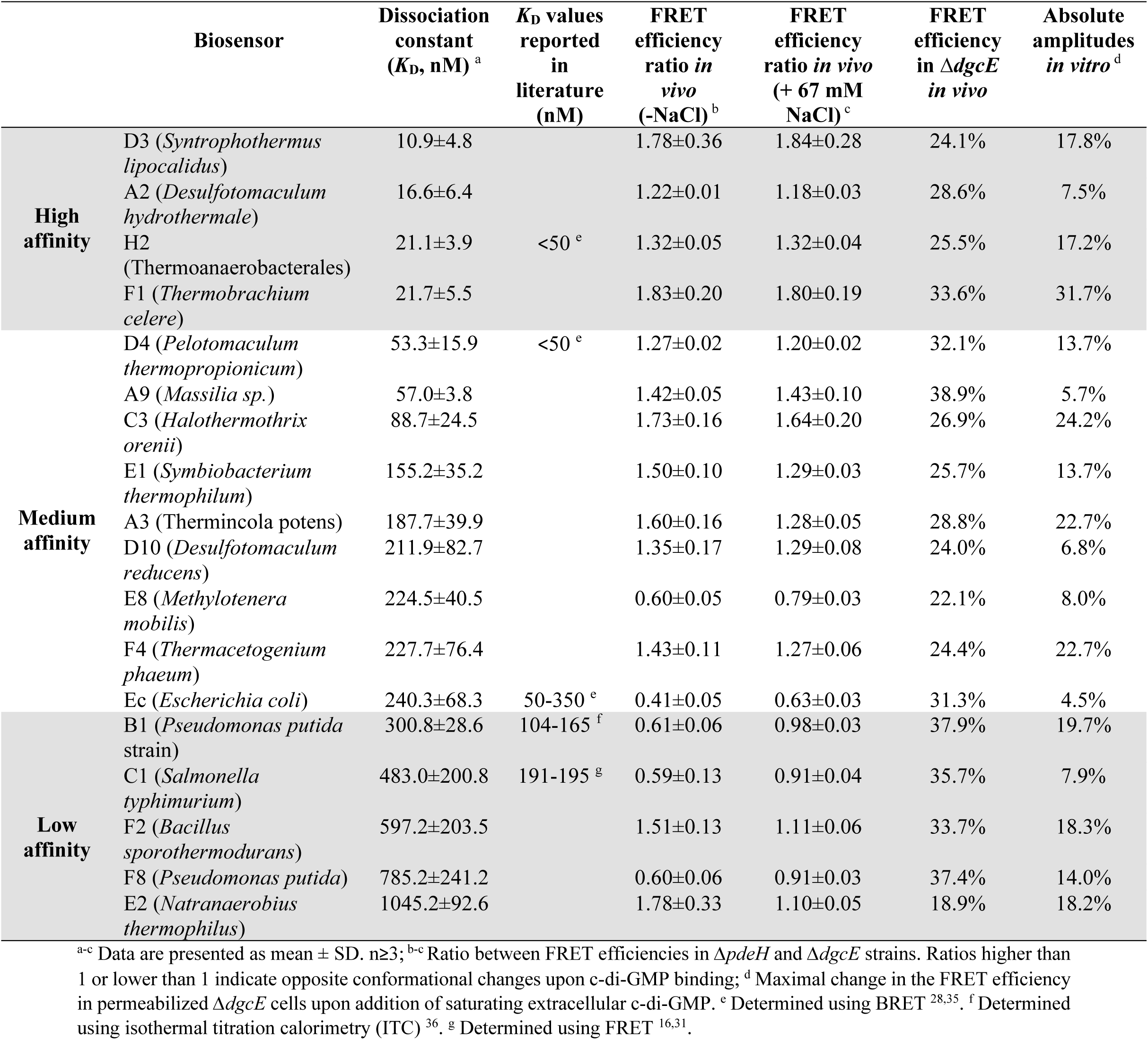
The toolbox of FRET-based c-di-GMP biosensors.

To assess the *in-vivo* affinity of different biosensors, we next compared their FRET efficiency in intact wildtype, *ΔpdeH* and *ΔdgcE* cells. We reasoned that a similarity between wildtype and *ΔpdeH* FRET values reflects greater occupancy of a particular sensor in wildtype cells, and thus its higher affinity for c-di-GMP (Fig. 2c and Supplementary Fig. 3a, b). Indeed, we observed a clear correlation between this *in-vivo* occupancy and the affinity of the sensor determined in permeabilized cells, confirming that the *K*_D_ values determined in permeabilized cells are consistent with the sensor affinity *in vivo*.

We further validated the dose-responses measured using flow cytometry with ratiometric microscopy measurements of FRET in a flow chamber ^37,38^, where permeabilized *ΔdgcE* cells were exposed to step-like stimulation by different concentrations of c-di-GMP (Fig. 3d). In contrast to flow cytometry, where each point in the c-di-GMP-binding curve was measured at an equilibrium state, this microscopy measurement of FRET allows to observe both dose-dependence and kinetics of the biosensor responses. Indeed, all three tested sensors with different affinities responded in the concentration range expected based on their *K*_D_ values. While the kinetics of the c-di-GMP binding was rapid (within 60 seconds) for all sensors, the apparent dissociation rate was much lower for the high-affinity F1 biosensor. Thus, low biosensor *K*_D_ is primarily due to the low off-rate of bound c-di-GMP, pointing to a tradeoff between affinity and time resolution of c-di-GMP measurements using FRET sensors.

**Fig. 3.**
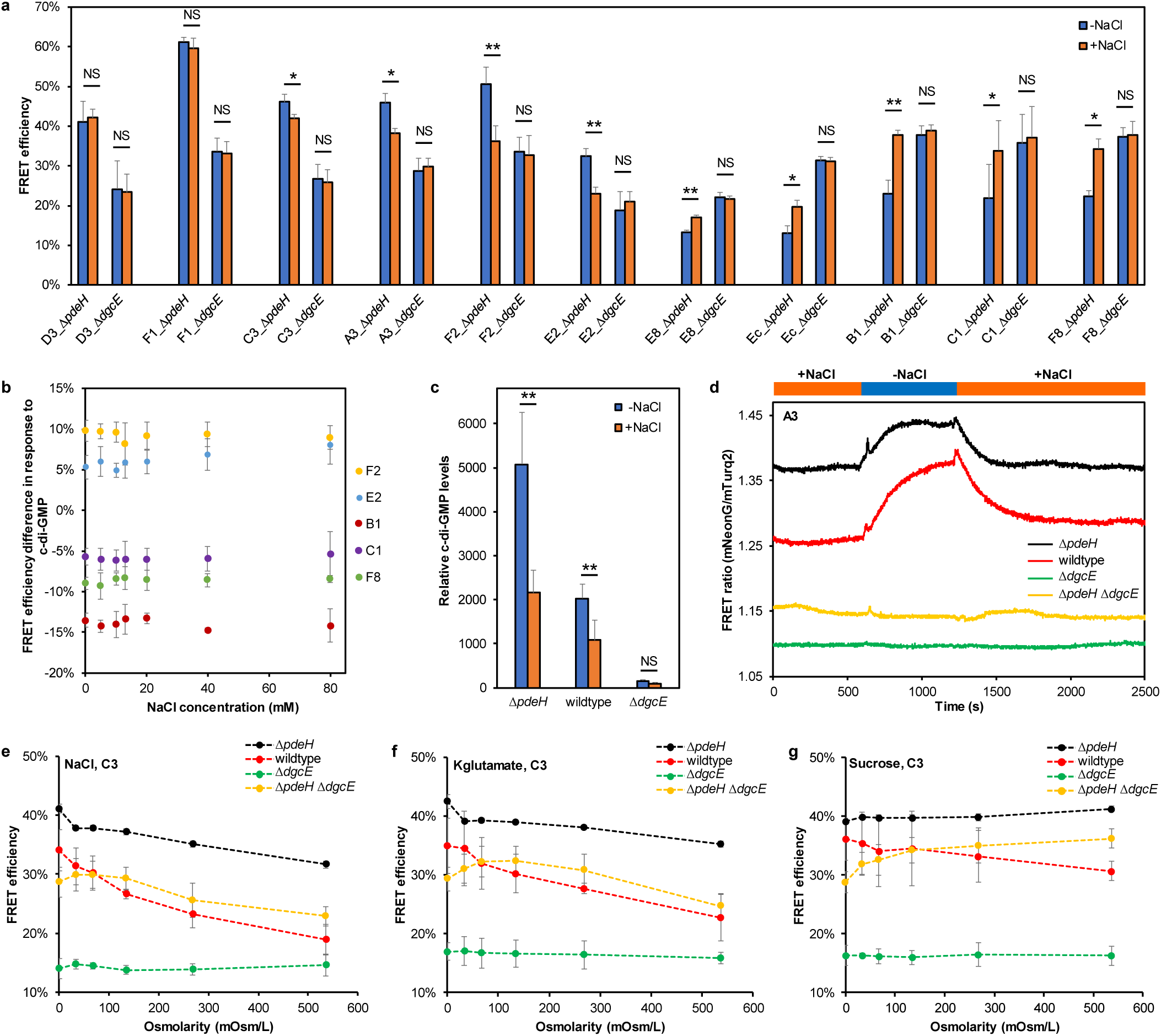
Effect of elevated NaCl concentration on the intracellular level of c-di-GMP in *E. coli*. **a,** FRET efficiency of indicated biosensors in *ΔpdeH* and in *ΔdgcE* strains, either in the absence of NaCl (-NaCl) or in presence of 67 mM NaCl (+NaCl). \*\**P* <0.01; \**P* <0.05; NS, no significant difference. *P* values were calculated using paired two-tailed *t-test*, n=3. Data are presented as mean ± SD. **b,** Difference in the FRET efficiency between exposure to high (810 nM) and low (1 nM) levels of c-di-GMP, for indicated biosensors in permeabilized *ΔdgcE* cells, at indicated concentrations of NaCl in the buffer. n=3. Data are presented as mean ± SD. **c,** The effect of NaCl on the intracellular c-di-GMP level in indicated *E. coli* strains, measured by LC-MS/MS. \*\**P* <0.01; NS, no significant difference. *P* values were calculated using paired two-tailed *t-test*, n≥3. Data are presented as mean ± SD. **d,** Ratiometric measurement of the response of A3 biosensor *in vivo* to the removal and addition of NaCl in the flow chamber, as indicated. Increased FRET ratio for this sensor indicates elevated c-di-GMP levels. **e-g,** Effects of indicated levels of ionic (sodium chloride and potassium glutamate) and non-ionic (sucrose) osmolytes on the FRET efficiency of C3 biosensor in intact cells of indicated strains. Higher FRET efficiency for this sensor indicates higher c-di-GMP levels. n≥3. Data are presented as mean ± SD.

### NaCl-dependent reduction of c-di-GMP level in *E. coli*

As mentioned above, the presence of a moderate concentration of NaCl (67 mM) in the buffer used for flow cytometry reduced the difference in FRET between *ΔpdeH* and *ΔdgcE* strains for most biosensors (Fig. 2a). To better understand the origin of this effect, we compared the values of FRET efficiency in *ΔpdeH* and *ΔdgcE* strains for 11 biosensors with top-ranked signal changes, in the absence and presence of NaCl (Fig. 3a). While all biosensors in *ΔdgcE* cells maintained the same level of FRET regardless of the presence of NaCl, the effect of NaCl on FRET in *ΔpdeH* cells varied among these biosensors (Fig. 3a and Table 1). There was a clear correlation between the magnitude of the NaCl impact on a biosensor in *ΔpdeH* cells and its *K*_D_ value, implying that biosensors with low affinity are stronger affected by NaCl (Supplementary Fig. 4).

We first hypothesized that NaCl may directly weaken sensor’s ability to change conformation upon c-di-GMP binding or reduce sensor’s affinity for c-di-GMP. To test this hypothesis, we measured the difference in FRET for 5 biosensors between low (1 nM) and high (810 nM) c-di-GMP in permeabilized cells, at concentrations of NaCl ranging from 0 to 80 mM (Fig. 3b). This covers the range of intracellular Na^+^ concentrations reported for live *E. coli* cells (5 to ∼13 mM) at extracellular NaCl ranging from 1 to 67 mM ^39^. However, no apparent impact of NaCl on *in-vitro* measured FRET could be observed. Consistently, NaCl does not directly affect sensors’ affinities measured in permeabilized cells (Extended Data Fig. 3).

Instead, extracellular NaCl might decrease the intracellular level of c-di-GMP in *ΔpdeH* (and wildtype) cells, thereby reducing the difference in c-di-GMP levels between these strains and *ΔdgcE.* A direct measurement of the cellular c-di-GMP levels in *E. coli* confirmed their decrease in *ΔpdeH* and wildtype cells upon incubation with NaCl (Fig. 3c). This could explain why the effect of NaCl was more pronounced for the low-affinity biosensors, which would be more sensitive to the reduction of c-di-GMP from high to intermediate levels in *ΔpdeH* cells, whereas high-affinity biosensors would remain saturated with c-di-GMP. Consistent with such modulation of cellular c-di-GMP levels by NaCl, a medium-affinity biosensor (A3; increased FRET ratio indicating elevated c-di-GMP levels) could rapidly respond to the removal and addition of NaCl in *ΔpdeH* or wildtype cells, but not in *ΔdgcE* cells (Fig. 3d). The lack of response in the *ΔdgcE* background does not seem to be a mere consequence of very low levels of the cellular c-di-GMP, since no response could also be observed in the *ΔpdeH ΔdgcE* strain that lacks both opposing enzymes and thus has higher level of c-di-GMP compared to *ΔdgcE* strain (Fig. 3d). Thus, the effect of NaCl on the c-di-GMP level appears to be mediated by DgcE.

To next test whether the decrease in c-di-GMP level is due to the increased osmolarity, we compared the effects of NaCl to those of other ionic and non-ionic osmolytes. Whereas c-di-GMP levels in the wildtype and *ΔpdeH* cells dropped with the increase of NaCl, KCl or potassium glutamate (Kglutamate) concentration (Fig. 3e,f and Extended Data Fig. 4a), the impact of non-ionic osmolytes at the same osmolarity was much weaker (Fig. 4g and Extended Data Fig. 4b). Therefore, the increase of osmolarity makes only minor contribution to the overall decrease of c-di-GMP, whereas ionic strength seems to have a major impact that is similar for Na^+^ or K^+^ ions.

**Fig. 4.**
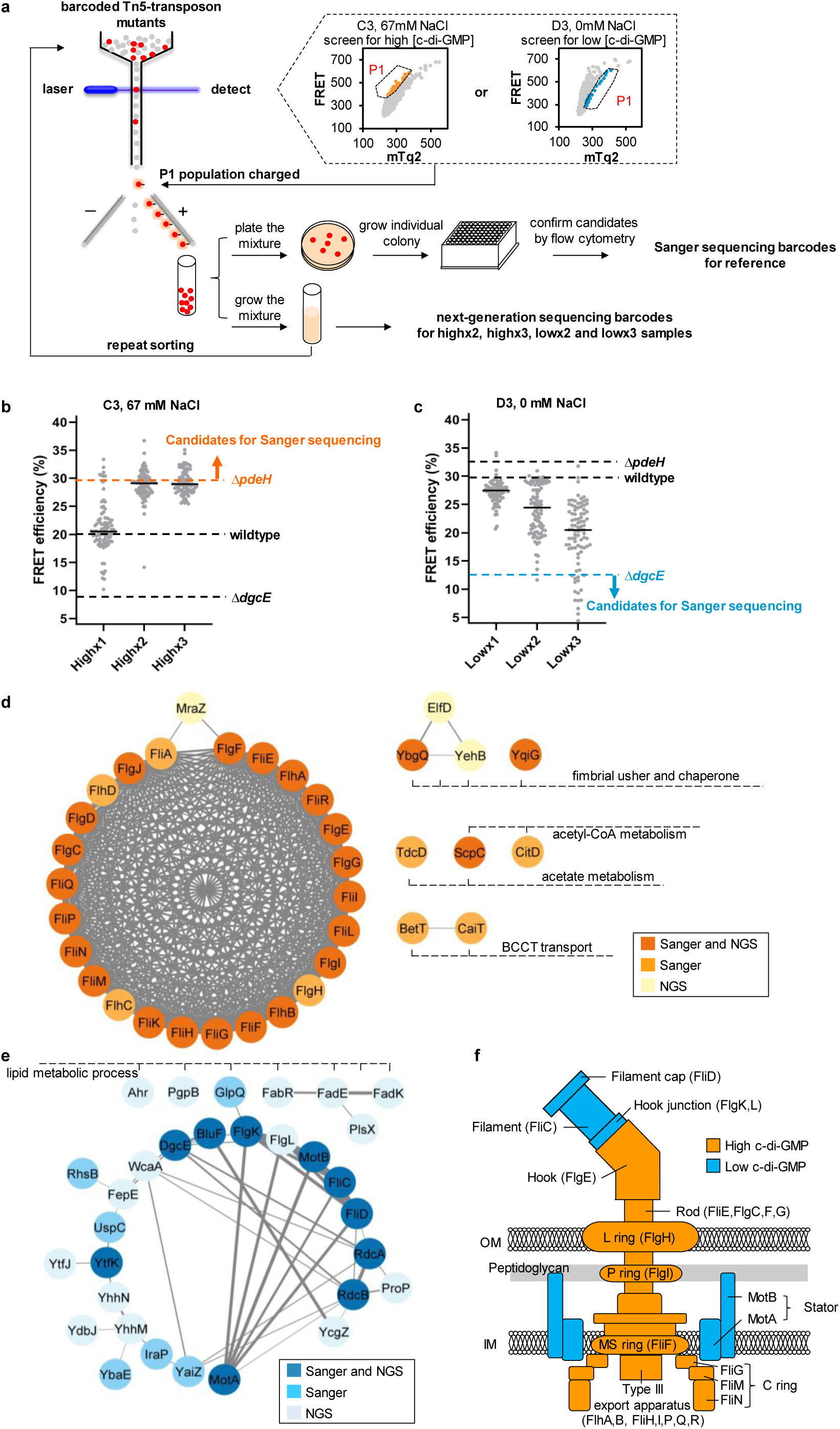
FRET-To-Sort application of the biosensor toolbox to identify genes that regulate c-di-GMP production. **a,** Schematics of FRET-To-Sort, based on selection and subsequent identification of mutants from a randomly barcoded transposon library through cell sorting based on the FRET signal. **b,** Distributions of FRET values for 93 colonies of mutants after the first (x1), second (x2) and third (x3) sorting, respectively, measured using flow cytometry. The C3 biosensor (*K*_D_ ∼89 nM) was utilized to sort out mutants with elevated c-di-GMP levels. The sorting buffer contained 67 mM NaCl to generally lower c-di-GMP levels and thus to increase selectivity for high-c-di-GMP sorting. Mutants confirmed to have higher c-di-GMP levels than *ΔpdeH* were genotyped by Sanger sequencing. **c,** Distributions of FRET values for 93 colonies of mutants after the first (x1), second (x2) and third (x3) sorting, respectively, measured using flow cytometry. The D3 biosensor (*K*_D_ ∼11 nM) was utilized to sort out mutants with low c-di-GMP levels. The sorting buffer contained no NaCl, to increase selectivity for low-c-di-GMP sorting. Mutants confirmed to have lower c-di-GMP levels than *ΔdgcE* were genotyped by Sanger sequencing. **d,e,** The networks and functional annotations of enriched mutants from Sanger sequencing and NGS data. Only networks with more than three proteins and significant functional enrichment annotated using DAVID ^68^ (*P* ≤ 0.05), are shown. Mutants selected for high c-di-GMP levels (**d,** orange) and for low c-di-GMP levels (**e,** blue) are shown. The networks were generated using the STRING database version 11.5 and visualized using Cytoscape 3.9.1. The thicker edge width of the links means higher confidence of the proposed functional association between two proteins. **f,** Effects of mutations in structural flagellar genes on c-di-GMP levels, with orange indicating up- and blue indicating down-regulation.

### FRET-To-Sort unveils network of genes affecting c-di-GMP levels

The c-di-GMP regulatory networks in bacteria are highly complex, and remain only partially understood even in model organisms such as *E. coli* ^40^. We envisioned that utilizing our biosensors that provide direct readout of cellular c-di-GMP levels, combined with transposon mutagenesis and flow-cytometry based measurement of FRET coupled to cell sorting, could provide a powerful tool (named FRET-To-Sort) to map the genome-wide regulation of c-di-GMP.

For that, we utilized a recently established *E. coli* library created by the Tn5-based transposon mutagenesis with random barcodes for transposon-site sequencing (RB-TnSeq) ^41,42^. This mutant pool was transformed with either medium-affinity C3 biosensor (*K*_D_ value ∼89 nM) to select mutants with high levels of c-di-GMP, or with high-affinity D3 biosensor (*K*_D_ value ∼11 nM) to select mutants with low levels of c-di-GMP. This transformed library was subjected to three sequential cycles of fluorescence activated cell sorting (FACS) (Fig. 4a). The stringency of sorting for mutations that upregulate c-di-GMP levels was further enhanced by adding 67 mM NaCl, which lowers average c-di-GMP level in *E. coli*, whereas the sorting for low c-di-GMP mutants was conducted without NaCl (Fig. 4a). When the enriched mutants obtained from each cycle of sorting were individually grown and analyzed using flow cytometry (Fig. 4a), the distribution of FRET efficiencies in the sorted population progressively shifted towards higher or lower c-di-GMP levels, respectively (Fig. 4b,c), validating our selection protocol. We then performed next-generation sequencing (NGS) on samples after two (x2) and three (x3) sorting cycles. For comparison, the colonies with confirmed high or low c-di-GMP levels (those above the *ΔpdeH* or below the *ΔdgcE* controls in Fig. 4b,c) were individually genotyped by Sanger sequencing of barcodes ^41^ (Supplementary Tables 2 and 3). Adding 134 mM NaCl to further lower basal level of c-di-GMP enabled sorting for high c-di-GMP mutants with the high-affinity D3 biosensor (Supplementary Fig. 5a). This produced the second set of high c-di-GMP mutants using Sanger sequencing (Supplementary Table 4), which showed similar results with those using C3 biosensor (Supplementary Fig. 5b).

When the enrichment of mutations was compared between high and low sorting samples in the NGS analysis for the second and third rounds of sorting, many genes showed reproducible differences (Extended Data Fig. 5, Extended Data Fig. 6, and Supplementary Tables 5 and 6). There was also a substantial overlap between Sanger sequencing and NGS results, which was particularly pronounced for high c-di-GMP mutants (Extended Data Fig. 6a,b). The overlap between samples was less pronounced for candidates with low c-di-GMP levels (Extended Data Fig. 6c), possibly because of reduced sensitivity of cell sorting at low FRET. Overall, a total of 92 genes were identified as significantly enriched in high c-di-GMP samples by NGS and Sanger (with 30 genes overlapping between C3 and D3 sensors), and 102 genes in low c-di-GMP samples (Supplementary Tables 2-6 and Table 2).

**Table 2.**
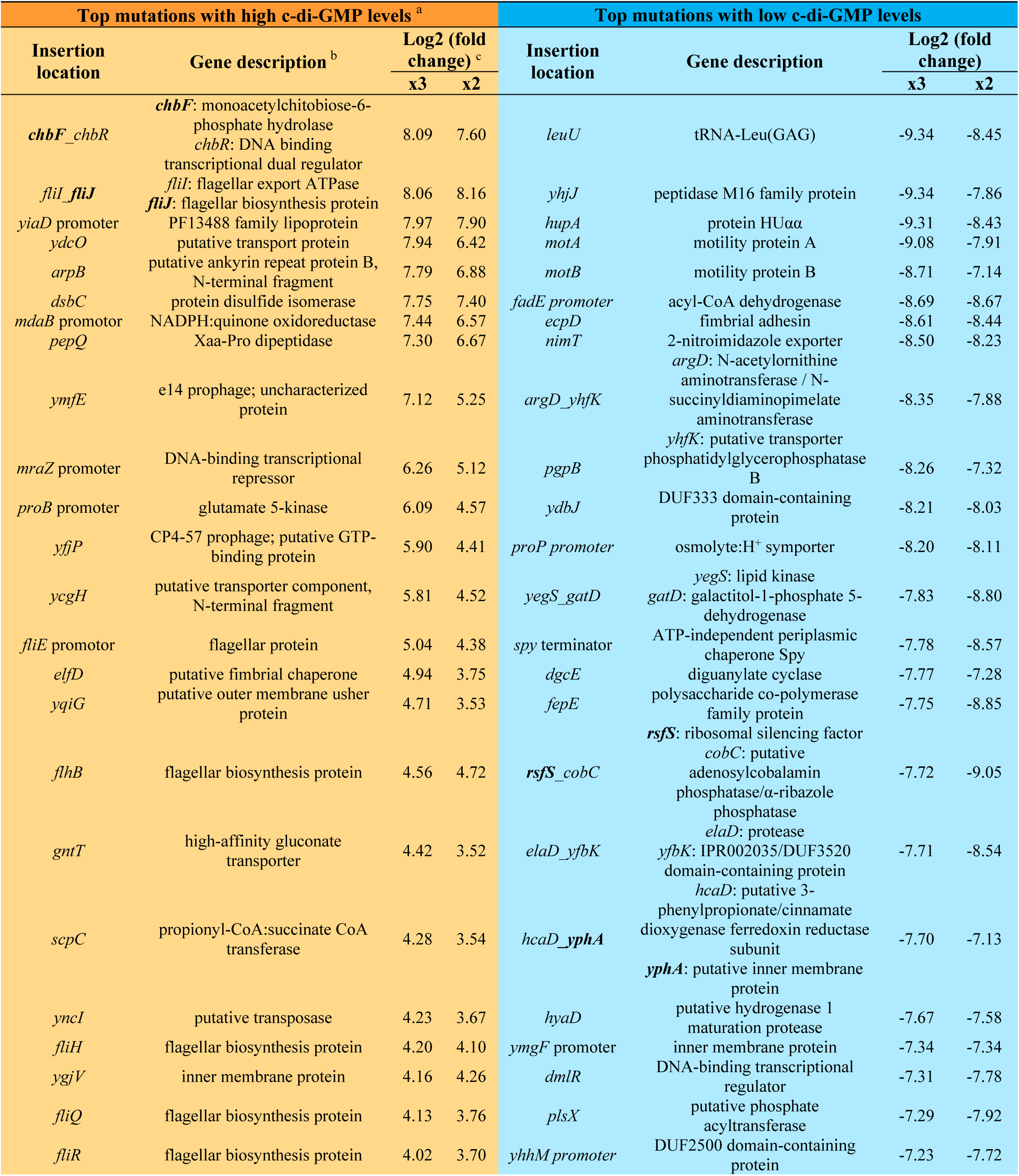
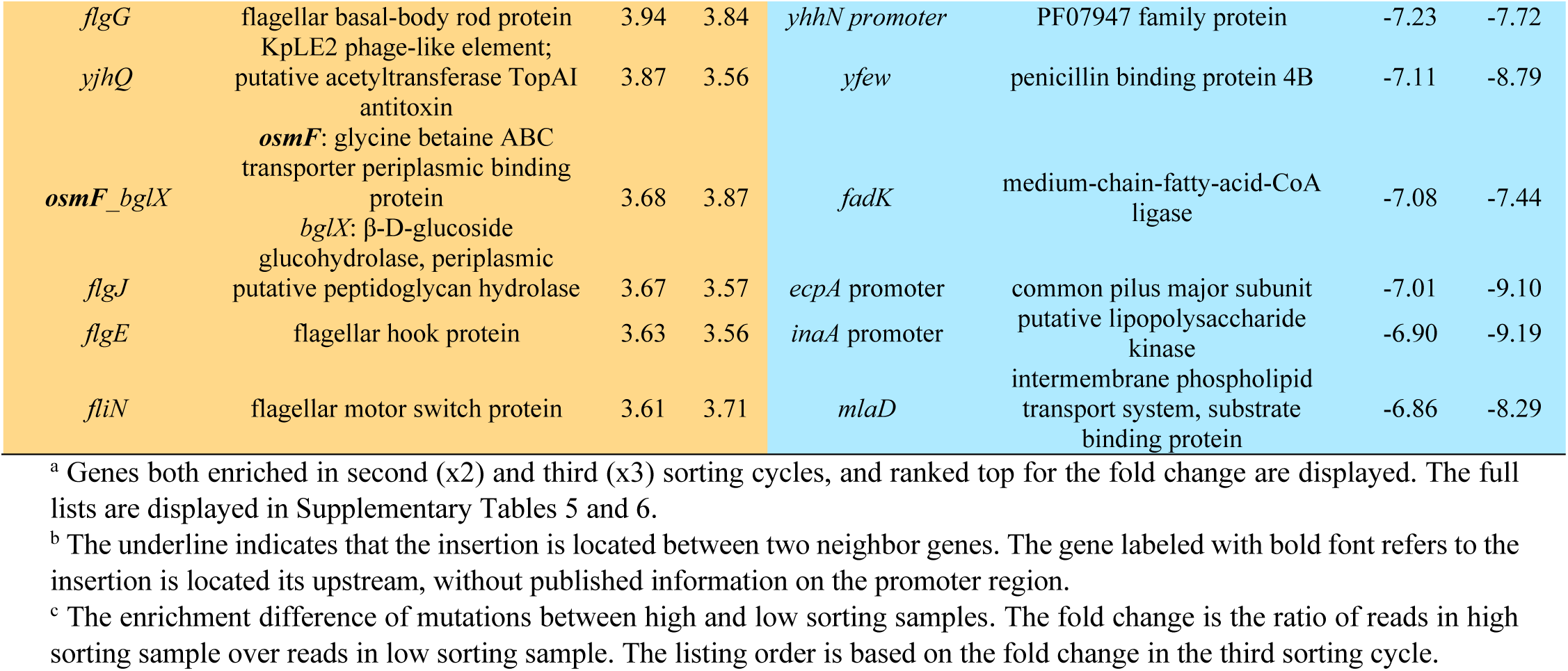
Mutations affecting c-di-GMP levels based on the NGS data.

We then generated networks of candidate genes obtained by NGS and Sanger sequencing through the STRING database (Fig. 4d,e), which integrate all known and predicted protein–protein associations ^43^. The most prominent group of mutations exhibiting high c-di-GMP levels mapped to flagellar genes (Fig. 4d). These include the class I genes encoding the master transcriptional activator of flagellar regulon FlhD/FlhC, as well as class II genes encoding components of the basal body and hook of *E. coli* flagellum and the sigma factor FliA ^44^. The latter is required for transcriptional activation of class III genes that include those encoding flagellin FliC, stator components MotA and MotB, and chemotaxis proteins, as well as PdeH. The activity of FliA is further known to be repressed in strains lacking structural class II genes ^44^, explaining similar phenotype of all class II gene mutations (see below). Also enriched was a mutation in the transcriptional regulator *yjjQ*, previously reported to repress multiple genes required for flagellar gene expression, capsule biosynthesis, outer membrane, stress resistance and biofilm formation in *E. coli*, including a cell-envelope stress responsive DGC, DgcN ^45,46^ . Other mutations showing elevated levels of c-di-GMP affected genes encoding fimbrial chaperone *elfD* and fimbrial usher proteins *ybgQ*, *yehB* and *yqiG*, outer membrane porins (*ompW*, *yiaD*), transcriptional regulator of cell wall genes (*mraZ*) or the periplasmic proteins (*dsbC*, *sanA*), suggesting that defects in the cell envelope may generally elevate c-di-GMP levels. Finally, several mutations with high c-di-GMP mapped to the acetate and acetyl-CoA metabolism.

Among mutants with reduced levels of c-di-GMP, we expectedly observed a prominent enrichment of genes encoding DgcE as well as its positive regulators RdcA and RdcB ^47^ (Fig. 4e). Also enriched was mutations in genes encoding the blue-light sensor BluF and its regulatory target YcgZ, which are known to impact biofilm formation and stress response ^48,49^. Other prominent groups were genes encoding proteins involved in lipid metabolism, including repressor of fatty acid biosynthesis FabR, and many genes associated with the stress response (Supplementary Tables 3 and 6), encoding for instance, the anti-adaptor protein IraP that stabilizes RpoS ^50^, the universal stress protein UspC and a stringent response modulator YtfK.

Surprisingly, mutations in class III flagellar genes, including *motA, motB, fliC*, were enriched among mutants with low c-di-GMP levels, in contrast to the impact of flagellar class II genes (Fig. 4f). As mentioned above, one possible mechanism connecting mutations in flagellar genes to the regulation of c-di-GMP could be via their impact on the expression of *pdeH*, which belongs to class III flagellar genes and similarly requires sigma factor FliA. Mutations in class II genes cause a defect in assembly of flagellar basal body and export apparatus, which results in the retention of FlgM, a negative regulator of FliA, that is normally secreted through the completed export apparatus to relieve repression of the class III genes ^44^. In order to test these hypotheses, we constructed deletion strains of several class II and class III genes in both the wildtype and *ΔflgM* backgrounds, as well as the deletion of the class I gene *flhD*. Consistent with our transposon mutant enrichment analysis, *ΔfliG* and *ΔfliH* cells lacking class II genes showed significantly increased c-di-GMP levels (Extended Data Fig. 7a,b). Deletion of *flhD* that is required for *fliA* and thus also *pdeH* expression increased the level of c-di-GMP, too, whereas deletion of *flgM* resulted in decreased c-di-GMP level. Importantly, no increase but rather even a mild decrease in c-di-GMP level was observed when *ΔfliG* and *ΔfliH* were deleted in *ΔflgM* background, suggesting that the positive effect of these mutations on c-di-GMP level is indeed mediated by FlgM.

Also consistent with our transposon mutagenesis, *ΔmotA* and *ΔfliC* cells showed low c-di-GMP levels, that were similar to the *ΔdgcE* control (Extended Data Fig. 7c). This reduction of c-di-GMP levels was significantly stronger compared to that caused by *flgM* deletion, and the effect of *ΔmotA* and *ΔfliC* mutations was still observed in *ΔflgM* background as well as in *ΔpdeH* background. Thus, the effect of class III gene mutations does not seem to be mediated by their effect on FlgM secretion and/or PdeH levels. Finally, we tested whether it might depend on DgcE. Since the level of c-di-GMP is already very low in *ΔdgcE*, we instead investigated the impact of *ΔmotA* and *ΔfliC* mutations in *ΔdgcE ΔpdeH* background with intermediate c-di-GMP levels. Also in this background, no significant effect of *ΔmotA* and *ΔfliC* could be observed, indicating that these mutations affect DgcE activity via a yet unknown mechanism (Extended Data Fig. 7d).

## Discussion

Signaling mediated by c-di-GMP is a key determinant of transition between planktonic and sessile lifestyles in many bacteria, but its complex dynamics and regulatory networks remain only partly understood. Here, we established a toolbox consisting of 18 c-di-GMP biosensors that enables monitoring intracellular c-di-GMP levels in living cells over a broad concentration range and with high temporal resolution. The flexibility of choosing biosensors with different, over two decades, affinities is important for applicability of our toolbox to a variety of bacteria and under diverse conditions, where cellular c-di-GMP level are known to vary widely ^1,2,20,21,25^.

To efficiently characterize this biosensor library in a high-throughput manner, in different bacterial strains and under various conditions, we established a flow-cytometry based FRET measurement assay. One application of this assay was to investigate the effects of osmolytes on c-di-GMP levels in *E. coli*. We observed that the presence of NaCl and other ionic osmolytes led to a decreased level of c-di-GMP, apparently dependent on DgcE, whereas non-ionic osmolytes had little effect. Although the nature of the underlying regulation remains to be elucidated, ionic osmolytes can depolarize the cell membrane of *E. coli* ^51^, providing potential connection to the activity of membrane cyclases. The negative impact of NaCl on c-di-GMP levels may be consistent with known inhibition of biofilm (curli) gene expression by NaCl ^52,53^, although this effect may be partly DgcE-independent ^12^. Notably, osmolarity- and/or salt-caused perturbation of cell envelope was also shown to affect c-di-GMP levels in other bacteria ^54,55^.

As another application of the flow-cytometry based FRET measurements, we established a FRET-To-Sort approach for the genome-wide mapping c-di-GMP regulatory networks. Here we utilized two biosensors with different affinities from our biosensor toolbox in combination with the recently constructed randomly barcoded Tn5 transposon *E. coli* mutant library ^42^ to enrich mutants exhibiting high or low c-di-GMP levels. FRET-To-Sort effectively confirmed genes with expected effects on c-di-GMP levels, including *dgcE* and its known positive regulators *rdcA* and *rdcB*. Moreover, also enriched were multiple genes known to affect the general stress response (e.g. *iraP, uspC*), stringent response (e.g. *ytfK*), and biofilm formation (e.g. *bluF*, *ycgZ*, *yjjQ*), consistent with the expected relation of these processes to c-di-GMP regulation in *E. coli* ^50,56^. In addition, we could identify a number of genes that had strong but previously not or little characterized impact on c-di-GMP signaling. Among mutations that caused elevation of c-di-GMP levels were those in components of Elf, Ybg, Yqi and Yeh fimbriae. Interestingly, these mutations were previously shown to cause defects in virulence ^57–59^, whereas a mutations in Ecp fimbria that rather lowered c-di-GMP levels was reported to enhance bacterial adhesion to host cells ^60^. Besides fimbrial genes, a group of mutations affecting outer membrane, periplasm or acetyl-CoA and lipid metabolism also caused changes in c-di-GMP levels, indicating that perturbation of the cell envelope and membrane composition might generally affect c-di-GMP signaling. Such cell-envelope stress is known to affect biofilm formation in *E. coli* ^61^ and in other bacteria ^62^, but how it leads to the c-di-GMP regulation in *E. coli*, and its possible relation to the observed impact of ionic osmolytes, remain to be elucidated.

The most prominent groups of genes affecting the c-di-GMP levels in *E. coli* were flagellar genes, with opposite effects observed for mutations in class I and II and in class III flagellar genes. The observed elevation of c-di-GMP in the absence of the master transcriptional activator FlhD-FlhC and of the sigma factor FliA, is in agreement with the established antagonism between expression of flagellar genes and c-di-GMP levels in *E. coli* ^1^. This negative regulation is primarily mediated by PdeH, which is expressed under control of FliA. Our results demonstrate that the FliA-dependent expression of *pdeH* can also explain the upregulation of the c-di-GMP level in class II *E. coli* mutants that are defective in the assembly of flagellar basal body and export apparatus. These defects preclude secretion of the anti-sigma factor FlgM, thus leading to inhibition of FliA activity and reduced PdeH expression. More generally, this regulation implies that any defect in the assembly of *E. coli* flagellar basal body will lead to the elevation of c-di-GMP, thus promoting transition to sessility. In contrast, inactivation of class III flagellar genes, including those encoding flagellin FliC and the stator components MotA and MotB, leads to strongly reduced levels of c-di-GMP, comparable to those observed in the absence of DgcE. We demonstrated that this effect was not dependent on FlgM or PdeH, but it is apparently mediated by the regulation of DgcE. These findings are generally consistent with previous observations that the functionality of the flagellar motor can affect the c-di-GMP production in different bacteria including *E. coli* ^21,63,64^. Such cross-regulation may be related to surface sensing (or mechanosensing) during early stages of transition towards the sessile lifestyle ^20,21^. Our genetic analysis suggest that DgcE could fulfill the function of sensing the flagellar motor integrity in *E. coli* through a yet unknown mechanism, which might, for example, rely on the perturbation of the membrane potential.

The FRET-To-Sort approach established here should be readily extended to investigate regulation of cellular levels of c-di-GMP in a variety of bacterial species where corresponding mutant libraries are available, including several important pathogens ^65,66^. More generally, it should be applicable to any other FRET-based second messenger or metabolite biosensors that are already available ^67^.

---

**Extended Data Fig. 1.**
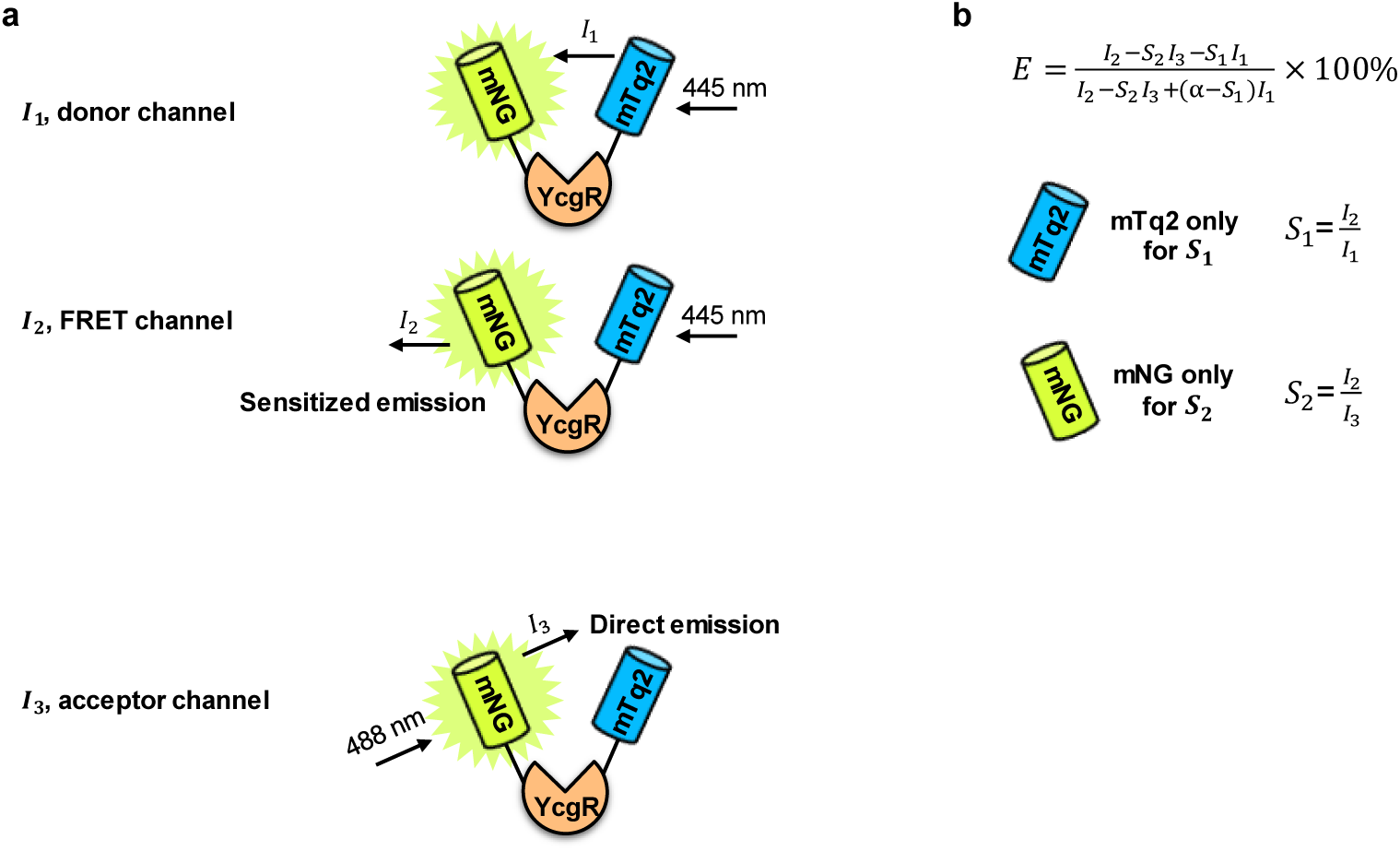
Illustration of the flow-cytometry based measurement of FRET. **a,** Three measured fluorescence signals include donor emission (*I_1_*) and sensitized emission of the acceptor (*I_2_*) upon direct donor excitation, and acceptor emission upon acceptor excitation (*I_3_*). **b,** Calculation of the FRET efficiency (*E*) from flow cytometry data. The mTq2 only and mNG only samples were used to calculate parameters *S_1_* and *S_2_*, respectively.

**Extended Data Fig. 2.**
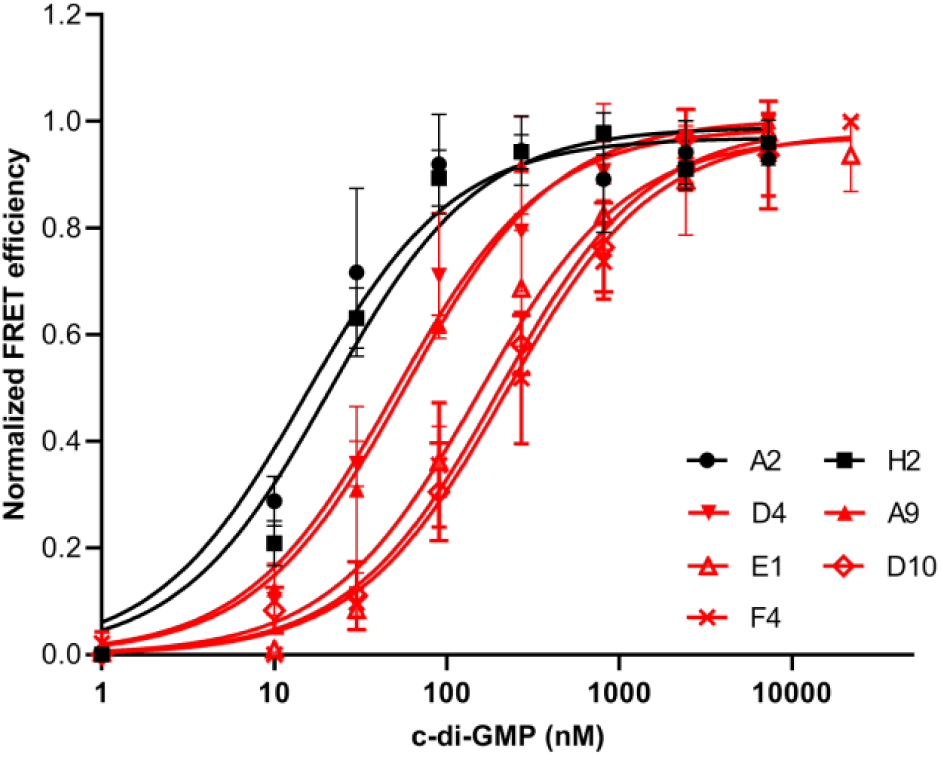
Dose responses of selected biosensors to c-di-GMP, measured in permeabilized *ΔdgcE* cells (i.e., *in vitro*). n=3. Data are presented as mean ± SD. Only biosensors chosen for the toolbox and not included in Fig. 2b are shown. Black or red color respectively indicates high or medium affinity.

**Extended Data Fig. 3.**
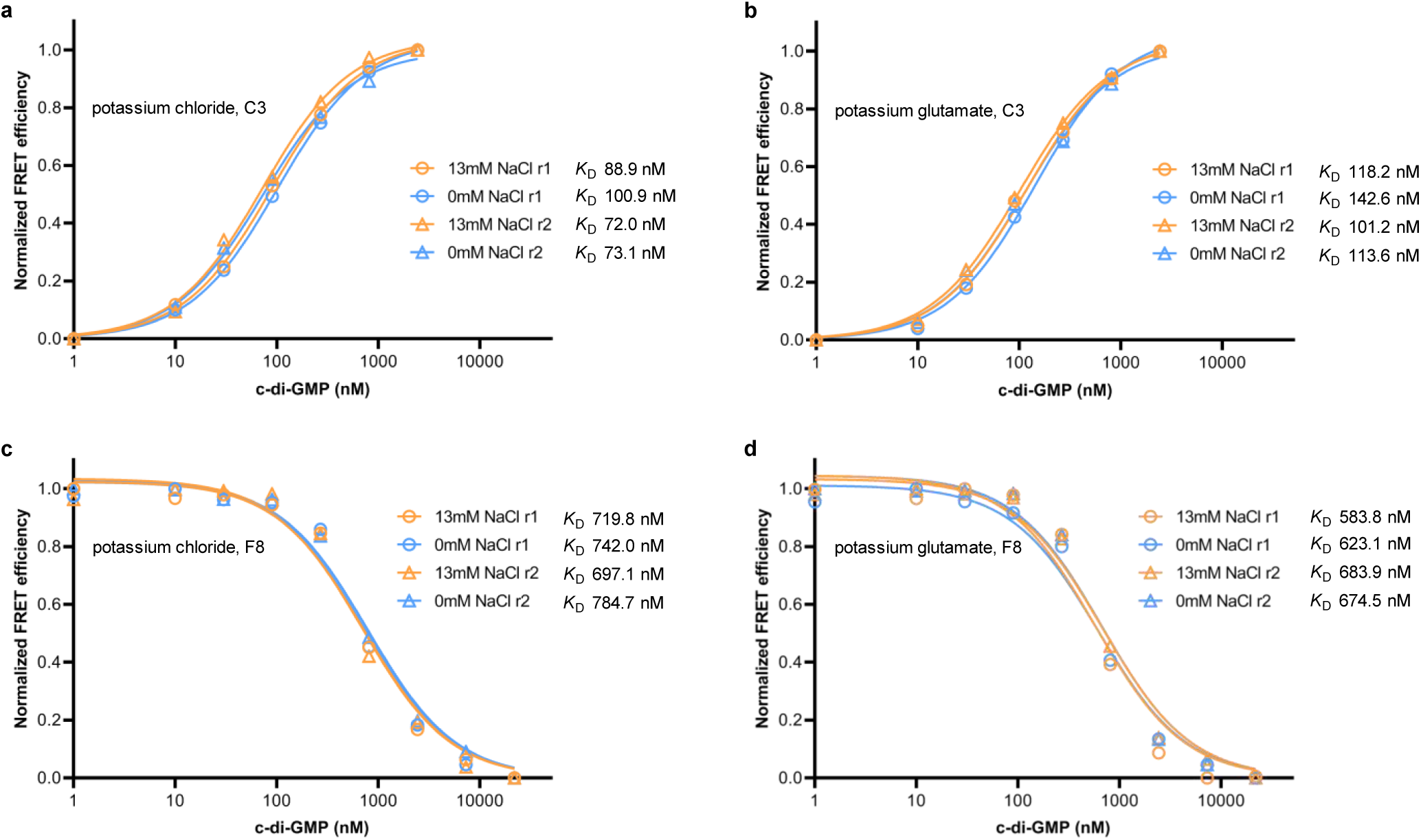
The affinity curves of C3 and F8 biosensors in the presence or absence of NaCl. Measurements were done in permeabilized *ΔdgcE* cells. The main source of potassium in the buffer is respectively potassium chloride (a and c) or potassium glutamate (b and d). The r1 and r2 represent independent biological replicate.

**Extended Data Fig. 4.**
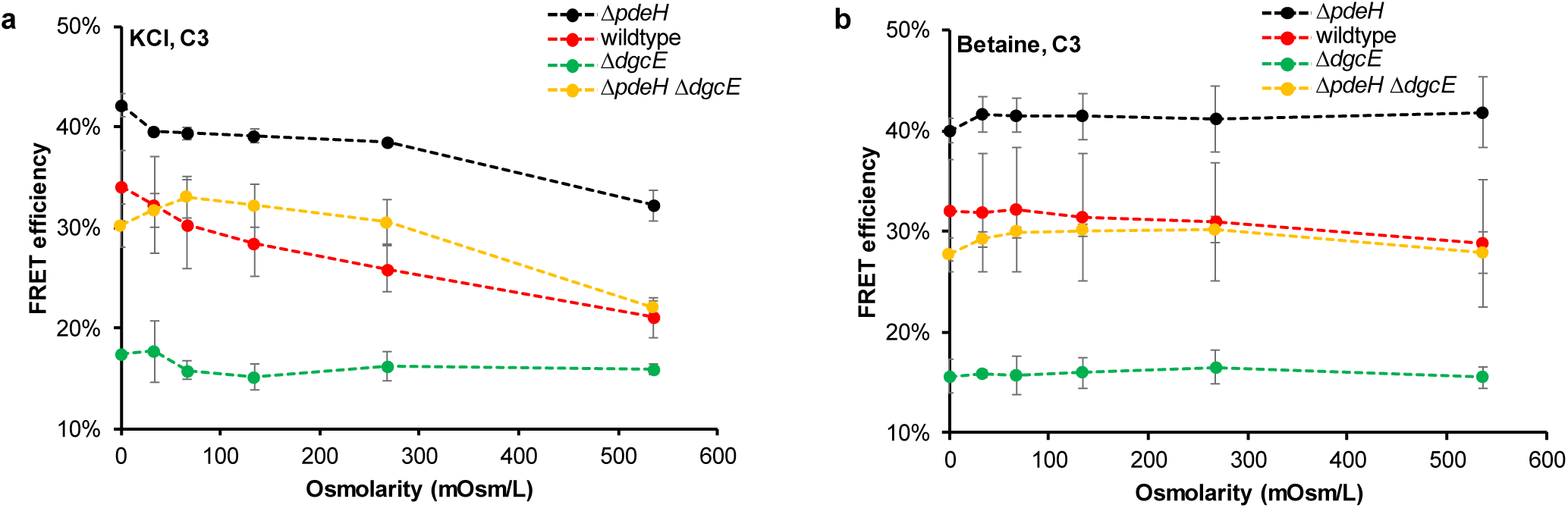
The effect of various levels of ionic (potassium chloride) and non-ionic (betaine) osmolytes on c-di-GMP. FRET efficiency of C3 biosensor was measured in live cells of indicated strains, in presence of indicated concentrations of potassium chloride (**a**) or betaine (**b**). Higher FRET efficiency indicates higher c-di-GMP levels. n≥3. Data are presented as mean ± SD.

**Extended Data Fig. 5.**
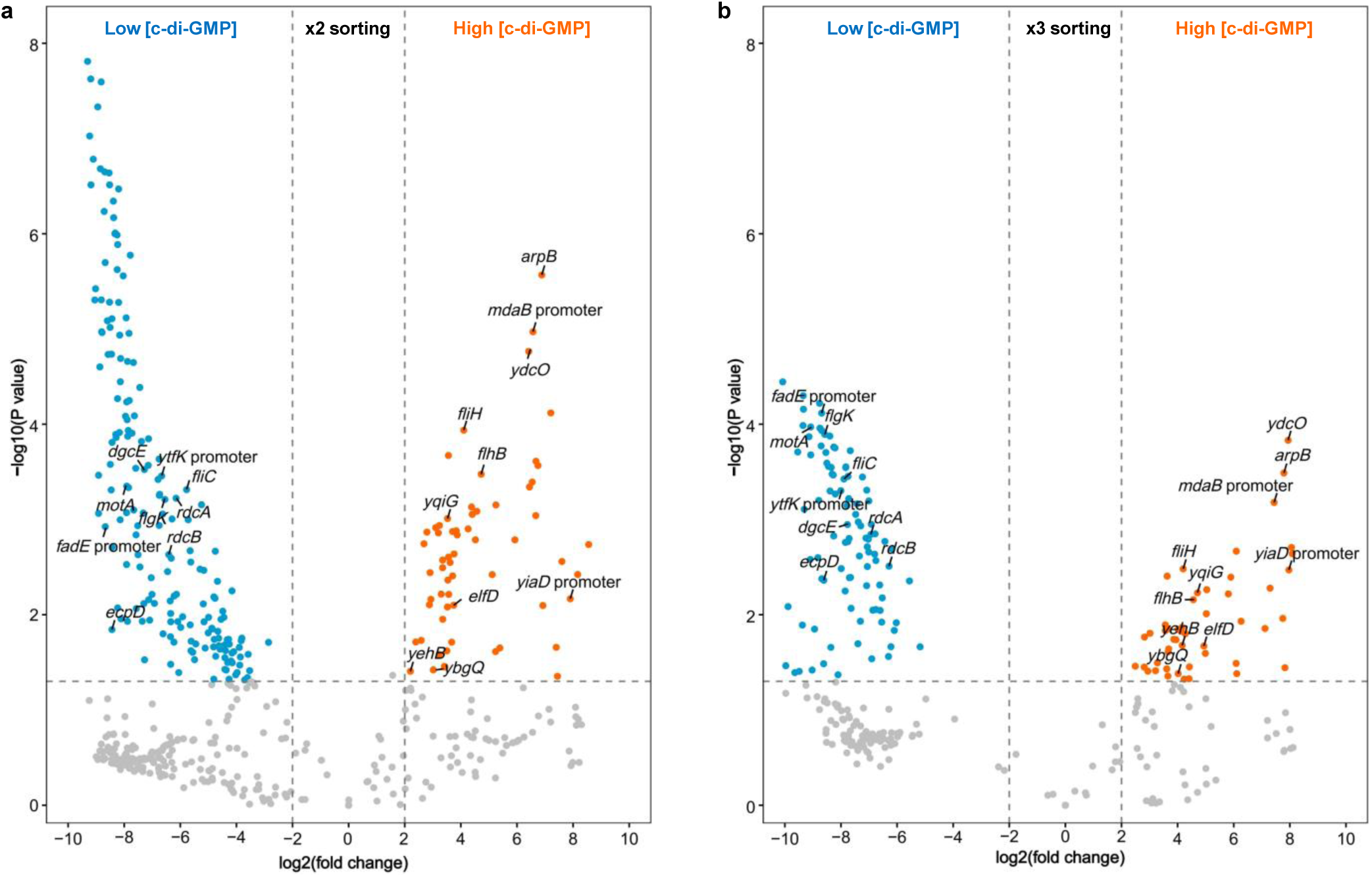
Volcano plots of NGS data show the enrichment difference between sorting for high and low c-di-GMP levels. **a,** Data for the second (x2) sorting cycle. **b,** Data for the third (x3) sorting cycle. n=2. Labeled are selected candidates enriched in both sorting cycles and shown in Fig. 4 or in Table 2.

**Extended Data Fig. 6.**
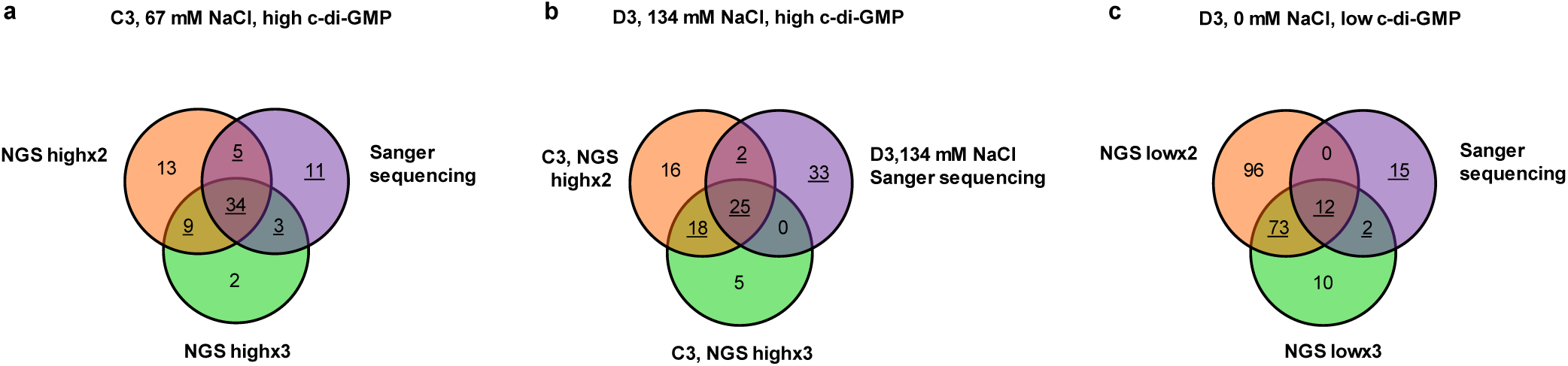
Venn Diagrams showing the overlap between mutations identified in Sanger sequencing data and NGS data. x2 and x3 indicate the second and the third sorting cycle, respectively. Underlined sets of mutants were chosen for further analysis. **a,** Sorting of mutants that elevate c-di-GMP levels using C3 biosensor (*K*_D_ ∼89 nM). The sorting buffer contained 67 mM NaCl to generally lower c-di-GMP levels and thus to increase selectivity for high-c-di-GMP sorting. **b,** Sorting of mutants that elevate c-di-GMP levels using D3 biosensor (*K*_D_ ∼11 nM) in the sorting buffer containing 134 mM NaCl, identified by Sanger sequencing, in comparison to the NGS data for the C3 biosensor from (a). **c,** Sorting of mutants that decrease c-di-GMP levels using D3 biosensor (*K*_D_ ∼11 nM). The sorting buffer contained no NaCl, to increase selectivity for low-c-di-GMP sorting.

**Extended Data Fig. 7.**
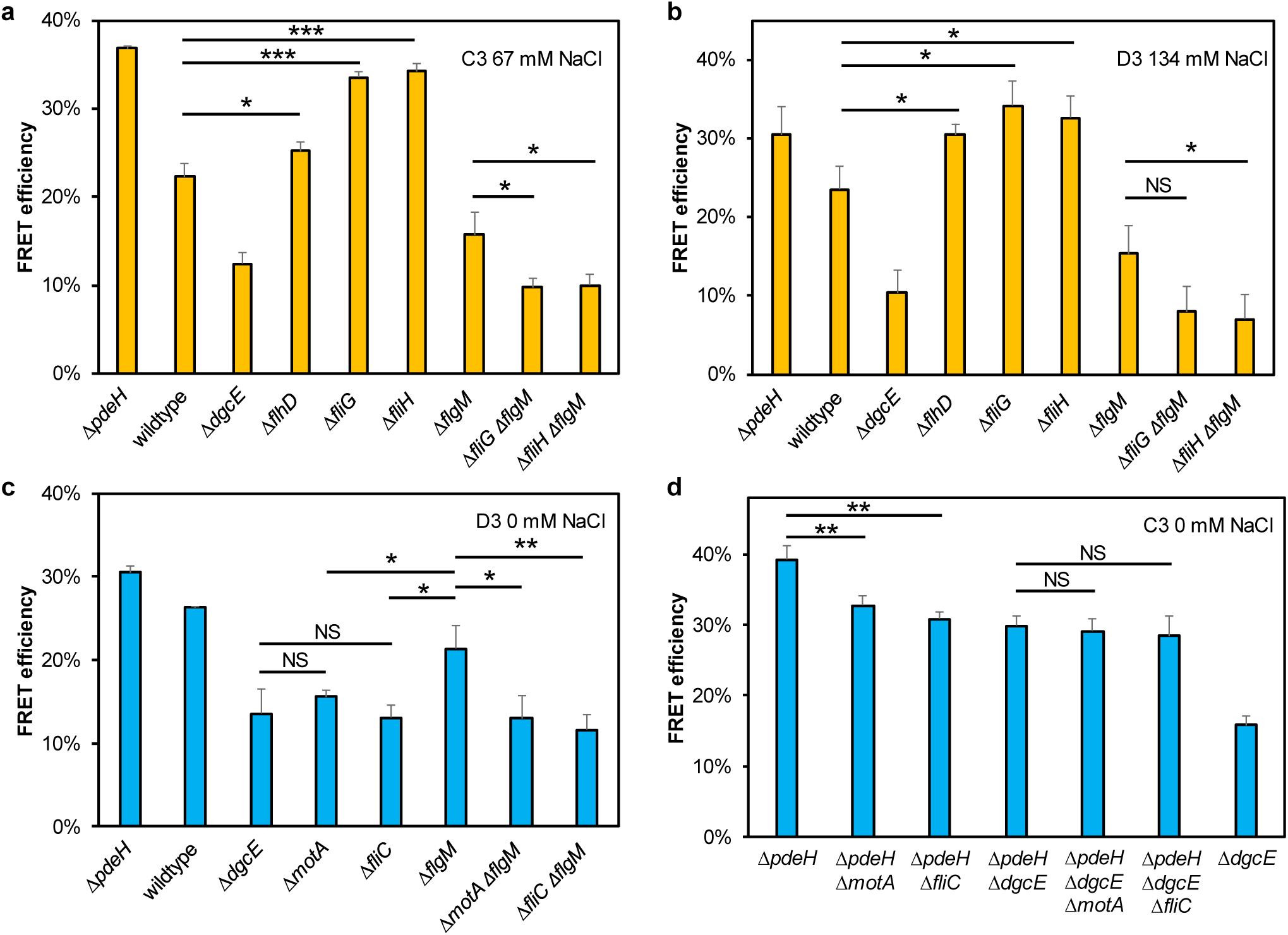
Effects of deletions in flagellar genes on the c-di-GMP levels in wildtype and mutant backgrounds. **a,b,** The impact of deletions of flagellar class I and II genes on c-di-GMP levels, detected using C3 biosensor in the presence of 67 mM NaCl (**a**) and D3 biosensor in the presence of 134 mM NaCl (**b**). **c,d,** The impact of deletions of flagellar class III genes on c-di-GMP levels, detected in the absence of NaCl, using D3 biosensor (**c**) and C3 biosensor (**d**). Higher FRET efficiency indicates higher c-di-GMP levels. \*\*\**P* <0.001; \*\**P* <0.01; \**P* <0.05; NS, no significant difference. *P* values were calculated using unpaired two-tailed *t-test*, n=3. Data are presented as mean ± SD.

## Materials and Methods

### Bacterial strains and plasmids

All strains and plasmids used in this study are listed in Table S7. *E. coli* strains were derived from the RpoS^+^ variant of W3110 ^69^, by PCR-product based inactivation of chromosomal genes ^70^ using plasmid pSIJ8 ^71^ to introduce gene deletions.

Library of genes encoding PilZ domain proteins ^28^ fused to mNeonGreen and mTurquoise2 was constructed and cloned into the pTrc99A vector using Gibson assembly ^72^. Following previous study ^28^, we used linkers AACGGCAGCCCATGG between mNeonGreen and PilZ domains, and GAGCTCTACAGGCTG between PilZ domains and mTurquoise2. The linear backbone was produced using restriction enzymes NcoI and HindIII, and insert fragments were amplified by PCR. *Ec ycgR* sequence was amplified from the *E. coli* W3110 genomic DNA, and the other 89 *ycgR* sequences were amplified from the published pET21/24-YNL-YcgR-91-mCherry plasmid library ^28^. For the construction of plasmids expressing mNeonGreen alone and mTurquoise2 alone, respective genes were separately cloned into the pTrc99A vector or into pBAD33 vector.

### Media and growth conditions

Cells were grown overnight in tryptone broth (TB; 1% tryptone, 0.5% NaCl) medium at 37°C with shaking at 200 rpm. Fifty microliters of overnight culture were diluted into 1 mL (96-deep well plates) or into 5 mL (24-deep well plates) fresh medium containing 20 μM isopropyl β-D-thiogalactoside (IPTG), for library screening or the *K*_D_ measurements, respectively. For control cultures carrying plasmids co-expressing mTurquoise2 and mNeonGreen from pBAD33 and pTrc99A vectors, respectively, the medium further contained 0.01% arabinose. Cells were harvested after 5 h of incubation at 37°C with shaking at 200 rpm. Bacterial culture was washed twice and resuspended in an appropriate buffer prior to measurements *E* values using flow cytometry or photobleaching microscopy. Where necessary, the culture medium was supplemented with antibiotics (100 μg/mL ampicillin or 34 μg/mL chloramphenicol).

### Measurements of FRET efficiency using acceptor photobleaching microscopy

Cells were grown as described above, and 1 mL culture was washed twice and resuspended in 70 μL tethering buffer. The surface of a glass bottom 96-well plate was treated with 0.1% poly-l-lysine for 15 min and rinsed with water twice. The cells were then added to the wells and incubated at room temperature for 10 min to allow attachment. Each well was gently rinsed with tethering buffer twice to remove unattached cells, and incubated with 200 μL tethering buffer afterward.

Measurements of the FRET efficiency by acceptor photobleaching were performed as described previously ^73,74^. Briefly, the measurement was conducted using a widefield Eclipse Ti-E inverted fluorescence microscope (Nikon, Japan) equipped with an X-Cite Exacte LED light source, a perfect focus system (PFS), and NIS-Elements AR software (version 4.40, Nikon). Excitation power was adjusted by controlling the fluorescence lamp power and neutral-density filters. Images were acquired with a 40x Plan Apo NA 0.95 objective, and recorded using iXon 897-X3 EMCCD camera (Andor) with EM gain set up to 250. The acquisition time was 1 s. The donor (mTurquoise2) was excited at 436/20 nm and its emission was detected at 472/30 nm. The acceptor (mNeonGreen) was excited at 504/12 nm and its emission was detected at 542/27 nm. The acceptor photobleaching was conducted by a 12-s excitation with 515 nm solid-state laser (CNI, Chuangchun, China). Two acceptor-channel and a hundred donor-channel images were taken before the photobleaching of the acceptor, and forty donor-channel and two acceptor-channel images were taken after the photobleaching.

FRET efficiency was calculated as follows:

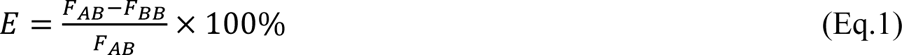

where background-corrected *F_AB_* and *F_BB_*were respectively donor fluorescence after and before acceptor photobleaching. Linear fitting of the donor fluorescence intensities versus time was performed for both pre- and postbleaching curves using RStudio to correct for donor photobleaching during data acquisition.

### Measurements of FRET efficiency using flow cytometry

The median fluorescence intensities of each channel were measured with BD LSRFortessa SORP cell analyzer (BD Biosciences, Germany). For each sample, a total of 30 thousand events was acquired. As illustrated in Extended Data Fig.1a, fluorescence intensities were measured in the donor channel (*I_1_*; 445 nm excitation, 470/15-nm emission), FRET channel (*I_2_*; 445 nm excitation, 520/35-nm emission) and acceptor channel (*I_3_*; 488 nm excitation, 520/30-nm emission). The power of both, 445 nm and 488 nm lasers was set at 75 mW. The median background level, obtained by measuring *ΔdgcE* strain carrying the empty pTrc99A vector, was always less than 2% of the signal in each channel and it was subtracted from each of the median fluorescence intensities.

The FRET efficiency was calculated from flow cytometric data following the previously established approach^33^ as follows (Extended Data Fig.1b):

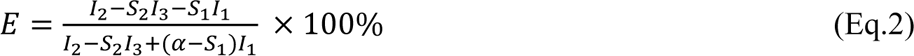

where *S_1_* corrects for the spectral bleed-through arising from the donor emission into the FRET channel, and *S_1_*equals to *I_2_/I_1_*, measured using a “donor-only” samples *i.e.* cells expressing mTurquoise2 alone*. S_2_* corrects for the bleed-though in the FRET channel arising from the not negligible direct excitation of the acceptor at 445 nm, and *S_2_* equals to *I_2_/ I_3_*, measured using “acceptor-only” samples, *i.e.* cells expressing mNeonGreen alone. Expression of mTurquoise2 and mNeonGreen in these control samples was in the same range as for the biosensors measured in this study.

The value of *α* is the calibration factor ^75^ relating the fluorescence intensity of an acceptor molecule in the *I_2_* channel to the intensity of a donor molecule in the *I_1_* channel for the mTurquoise2-mNeonGreen FRET pair for a given experimental setup. This value was determined as *α* = 0.739±0.081 in our case following the previously described approach ^33^, by calculating the ratio of the slope and the intercept from the plot of 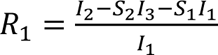, where 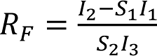 and 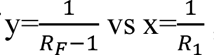. Notably, given the number of available biosensor constructs, we were able to obtain the *α* calibration factor from the interpolation of 48 values from independent one-to-one donor/acceptor samples having different FRET efficiencies (Supplementary Fig. 6).

For measurements of *E* values in intact cells, cells were grown as described above, washed twice and resuspended in tethering buffer (20 mM potassium phosphate, 0.1 mM EDTA, 1 µM methionine, 10 mM lactic acid, pH 7) with or without the addition of 67 mM NaCl for 1 h to equilibrate prior to measurements. The effect of osmolarity on c-di-GMP levels was tested using tethering buffer supplemented with different concentrations of osmolytes as indicated.

For measurements of *K*_D_, *ΔdgcE* cells carrying biosensors were grown as described above, washed twice and resuspended in the assay buffer (50 mM potassium phosphate [pH 7.4], 100 mM KCl, 0.5 mM MgCl_2_, 13 mM NaCl), of which the composition was similar to ion levels in live bacteria ^39,51^. After adding 1% toluene, cell suspension was incubated at 24°C with shaking at 300 rpm for 10 min, and diluted 1:20 into the assay buffer containing different concentrations of c-di-GMP. The mixture was vortexed and left at room temperature for at least 1 h to equilibrate before the measurement. The *K*_D_ values were calculated from the c-di-GMP dose-response curves using allosteric sigmoidal model in GraphPad Prism (version 9.0, Graphpad software Inc., San Diego, CA).

### Ratiometric FRET measurements using microscopy

The ratiometric FRET measurements were performed using the Eclipse Ti-E inverted fluorescence microscope (Nikon, Japan) as described previously ^76^. Cells were attached to coverslips at the bottom of a flow chamber, and simultaneously imaged in the donor (mTurquoise2, detected at 472/30 nm) channel and the acceptor (mNeonGreen, detected at 542/27 nm) channel using Optosplit (OptoSplit II, CAIRN Research). Images were acquired with a 40x Plan Apo NA 0.95 objective, and continuously recorded using iXon 897-X3 EMCCD camera (Andor) with the acquisition time of 1 s. The flow was controlled using a syringe pump (Harvard Apparatus). The same region of interest (ROI), fully covered with a monolayer of cells, was selected in each image throughout the entire time series. The FRET ratio between fluorescence emission in the acceptor and in the donor channel while exciting donor fluorescence was recorded.

To measure response kinetics of selected biosensors, the *ΔdgcE* cells carrying biosensors were grown and then permeabilized using toluene as described above. One milliliter of permeabilized cells in the assay buffer was washed twice and concentrated into a 30 μL volume. The coverslip was treated by 0.1% poly-l-lysine for 15 min and rinsed with water twice. Permeabilized *ΔdgcE* cells were added on the coated coverslip for 20 min and mounted onto the flow chamber. A stepwise addition and subsequent removal of indicated concentrations of c-di-GMP in the assay buffer was applied at a constant rate of 0.3 mL/min.

To measure impact of NaCl on c-di-GMP levels in live cells, *ΔpdeH*, wildtype, *ΔdgcE*, or *ΔpdeH ΔdgcE* strains carrying A3 biosensors were grown as described above, and 1 mL culture was washed twice and resuspended in 50 μL tethering buffer. To avoid the effect of poly-l-lysine on intracellular c-di-GMP levels, cells were added on coverslips without poly-l-lysine coatings. After attachment for 40 min, the coverslip was mounted onto the flow chamber. The tethering buffer with the addition and subsequent removal of 67 mM NaCl was applied at a constant rate of 0.3 mL/min. The cell detachment in ROIs was checked manually throughout the entire time series. We found that *E. coli* W3110 attached well on coverslips without poly-l-lysine coatings under the flow rate of 0.3 mL/min.

### Measurement of c-di-GMP levels using liquid chromatography with tandem mass spectrometry (LC-MS/MS)

To assess the relative c-di-GMP concentration changes due to the presence of NaCl, c-di-GMP levels were quantified by LC-MS/MS. The cells were grown as described above, and 1 mL of overnight culture was diluted into 100 mL fresh TB medium. Cells were harvested at desired optical density (OD_600_ ∼1.2) by spinning down 10 mL culture volume. The resulting pellet was resuspended in 50 mL tethering buffer with the presence or absence of 67 mM NaCl and stayed at room temperature for 1 h. Cells were harvested by centrifugation and the pellet was extracted on ice as follows.

The extraction was performed as previously described ^77^, slightly modified for the extraction of c-di-GMP. The extraction liquid was 50% (v/v) methanol in TE-buffer (10 mM TRIZMA, 1 mM EDTA, pH 7.0), stored at - 20°C. The cell pellet was resuspended in 150 μL of ice-cold extraction fluid and then mixed with 150 μL of ice-cold 100% chloroform in a precooled 1.5 mL Eppendorf tube. The mixture was incubated at 1°C for 2 h, with shaking in an Eppendorf ThermoMixer. Subsequently, the samples were centrifuged at -10°C for 10 min (12,000 g). The upper phase of the two-phase system was filtered (0.22 µm, PTFE, 4 mm diameter, Phenomenex, USA) and stored at -20°C. The analysis and quantification was performed as previously described ^77^, slightly modified for c-di-GMP ^78^.

### FRET-To-Sort

To enable FRET-based sorting of mutants according to their c-di-GMP levels, a selected biosensor from our toolbox was transformed into the previously described randomly barcoded Tn5 transposon mutant library ^42^ by electroporation. After 2 h of recovery at 37°C, the cells were diluted, plated in LB-agar supplemented with ampicillin and kanamycin, and grown overnight at 37°C. The colonies were washed off the plates using 20% (w/w) glycerol in TB medium, and stored at -80°C in 500 mL aliquots.

Ten microliters of the glycerol stock of the transformed Tn5 transposon mutant library were used to inoculate the culture grown overnight in 5 mL TB medium supplemented with ampicillin at 37°C with shaking at 200 rpm. Fifty microliters of overnight culture were diluted into 5 mL fresh medium containing 40 μM IPTG. Cells were incubated for 4 h at 37°C with shaking at 200 rpm. A hundred microliters of the culture were diluted in 2 mL tethering buffer in the presence or absence of NaCl as indicated. After 1 h of equilibrium, the cells were sorted at 4°C by the cell sorter (BD FACSAria^TM^ Fusion, BD Biosciences, Germany).

The cells were excited at 445 nm, and the donor emission (mTurquoise2, detected through 470/15-nm filter) and the FRET signal (acceptor emission, detected through 520/35-nm filter) were measured. Doublets and debris were excluded by gating of forward scatter (FSC) and side scatter (SSC). The cells with a high or low ratio between signals in the FRET channel and the donor emission channel were selected (1-2% of the population), as indicated in Fig.4a. A tube containing 2ml of TB medium supplemented with ampicillin was used to collect the sorted cells, with 50 thousand cells per tube.

The sorted cells were allowed to recover at 37°C for 2 h without shaking, and divided into two parts: 1). One milliliter of recovered samples was diluted, plated in LB-agar supplemented with ampicillin and kanamycin and grown overnight at 37°C. Single colonies were grown overnight, diluted at ratio of 1:100 with fresh medium containing 40 μM IPTG, grown for 4 h at 37°C with shaking at 200 rpm, resuspended in tethering buffer in the presence or absence of NaCl, and analyzed by flow cytometry for c-di-GMP levels, as described above. Those colonies showing either higher c-di-GMP levels than *ΔpdeH* strain or lower c-di-GMP levels than *ΔdgcE* strain were performed colony PCR using Q5 polymerase (New England Biolabs, Ipswich, Massachusetts) with U1 and U2 primers. The barcode sequence was obtained by Sanger sequencing. The other one milliliter of the recovered samples was added 4 mL of fresh TB media supplemented with ampicillin and grown overnight at 37°C with shaking at 200 rpm. Fifty microliters of overnight culture were grown in fresh TB medium containing 40 μM IPTG before repeating the sorting step. The rest of overnight culture was used to perform NGS as follows.

Genomic DNA was extracted using NucleoSpin Microbial DNA kit. Barcode sequences were amplified by PCR using Q5 polymerase with barcoded primers for NGS ^41^. The PCR reactions were pooled together (10 µL from each reaction), purified using 0.9X of AMPure beads (Beckman Coulter, Brea, California) and sequenced in the GeneCore facility (EMBL, Heidelberg, Germany). The samples were run using the Novaseq 6000 SP Reagent Kit (100 cycles, Illumina, San Diego, California) with 30% PhiX, providing us around 8 million reads per sample. The results of the sequencing were processed and analyzed as followings.

The barcodes of each sample were extracted using the CutAdapt tool ^79^, with the sequences U1 and U2 as adaptors, a minimum size of 20bp, and quality higher than 20. The barcodes were then reverse complemented, counted and matched with the Tn5 transposon mutant library map. The read of each gene is the sum of the frequency of barcodes inserted in the same gene, which was subsequently normalized by total reads of all genes in the sample. We used unsorted sample as a reference to filter data. The genes with an average read that was 2.5-fold higher than its corresponding unsorted reference were selected for further analysis. The significance of difference in gene enrichment between high and low sorting samples was analyzed using unpaired *t-test* in Cyber-T web server ^80^.

---

**Supplementary Fig. 1.**
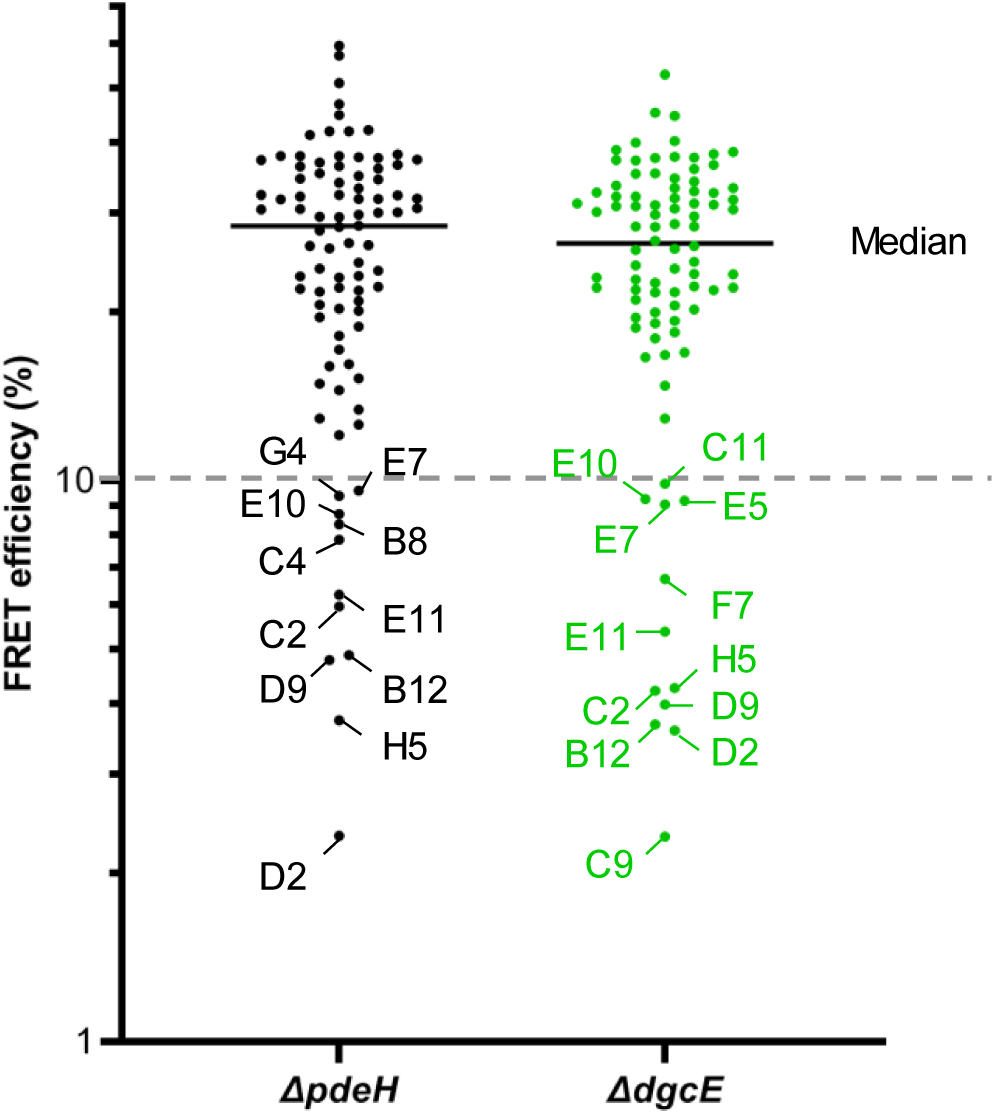
Distributions of FRET efficiency for all biosensors in *ΔpdeH* and in *ΔdgcE* backgrounds. Biosensors with FRET efficiency below 10% are labeled. The biosensors showing low values of FRET efficiency in both deletion strains were excluded for further analysis. The data showed the measurement conducted in the presence of 67 mM NaCl in tethering buffer using flow cytometry. Average values are presented.

**Supplementary Fig. 2.**
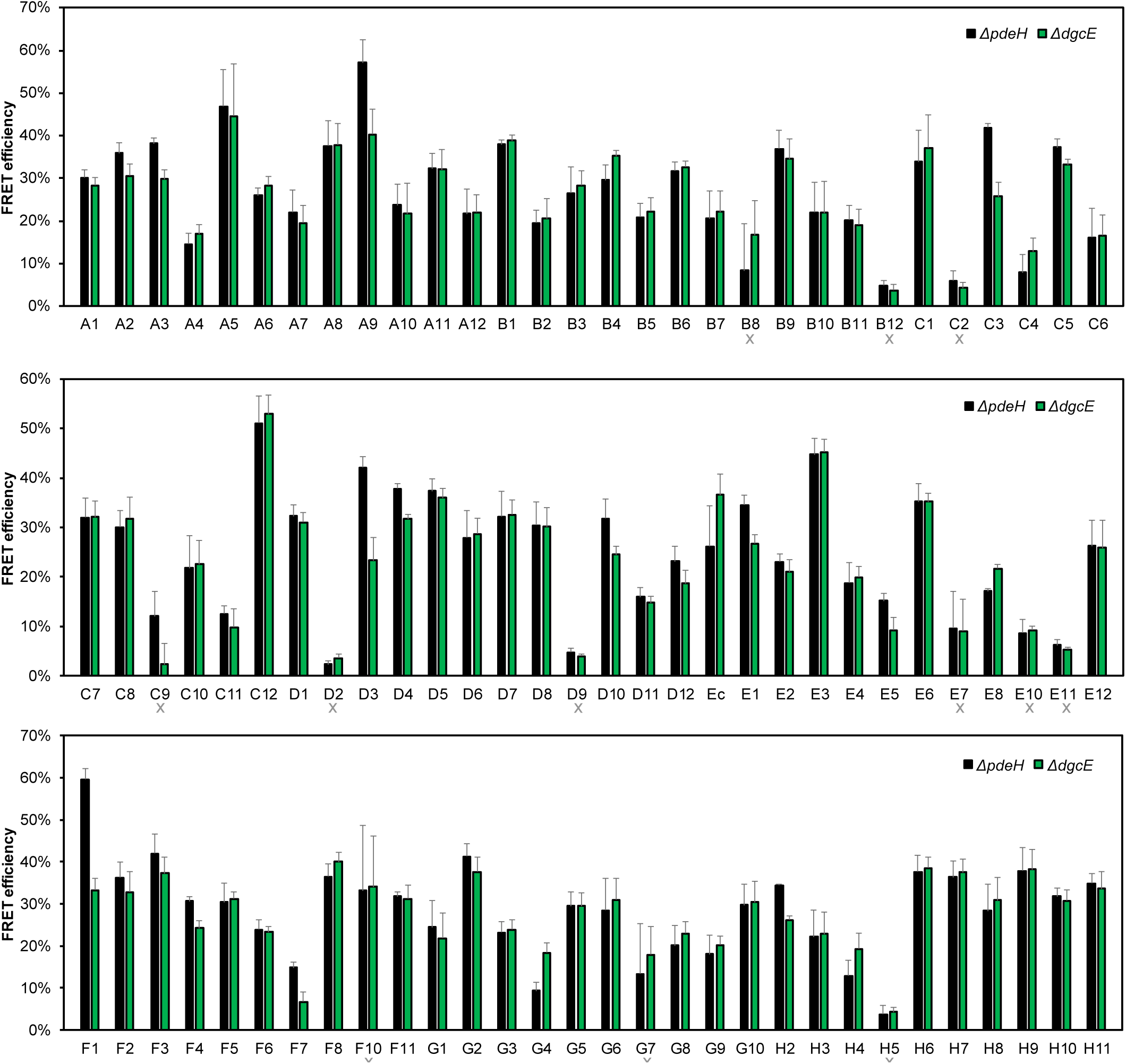
FRET efficiency of each individual biosensor in *ΔpdeH* and in *ΔdgcE* backgrounds. The data showed the measurement conducted in the presence of 67 mM NaCl in tethering buffer using flow cytometry. n=3. Data are presented as mean ± SD. The biosensors labelled with a grey cross underneath were excluded from further analysis due to low values of FRET efficiency (<10%) or high coefficient of variances in both deletion strains.

**Supplementary Fig. 3.**
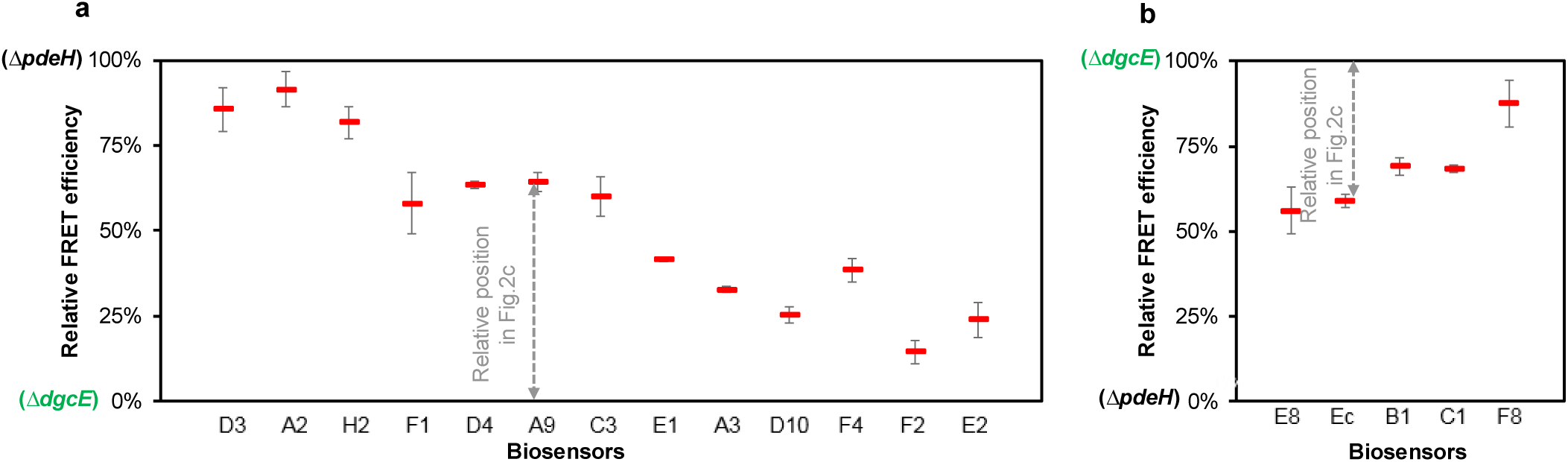
Illustrations explaining the position of the wildtype values relative to *ΔdgcE* and *ΔpdeH* values in Fig.2c. **a,** For biosensors showing higher FRET signals after binding c-di-GMP, the FRET efficiency in *ΔdgcE* and in *ΔpdeH* was assumed as 0 and 100%, respectively. **b,** For biosensors showing opposite conformation change, i.e. lower FRET signals upon binding c-di-GMP, the FRET efficiency in *ΔdgcE* and in *ΔpdeH* was assumed as 100% and 0, respectively. The relative FRET efficiency in wildtype is always located between 0 and 100%. The distance between *ΔdgcE* and wildtype was defined as the relative position, indicated using the grey dash arrow. The values were measured in the absence of NaCl. n=3. Data are presented as mean ± SEM.

**Supplementary Fig. 4.**
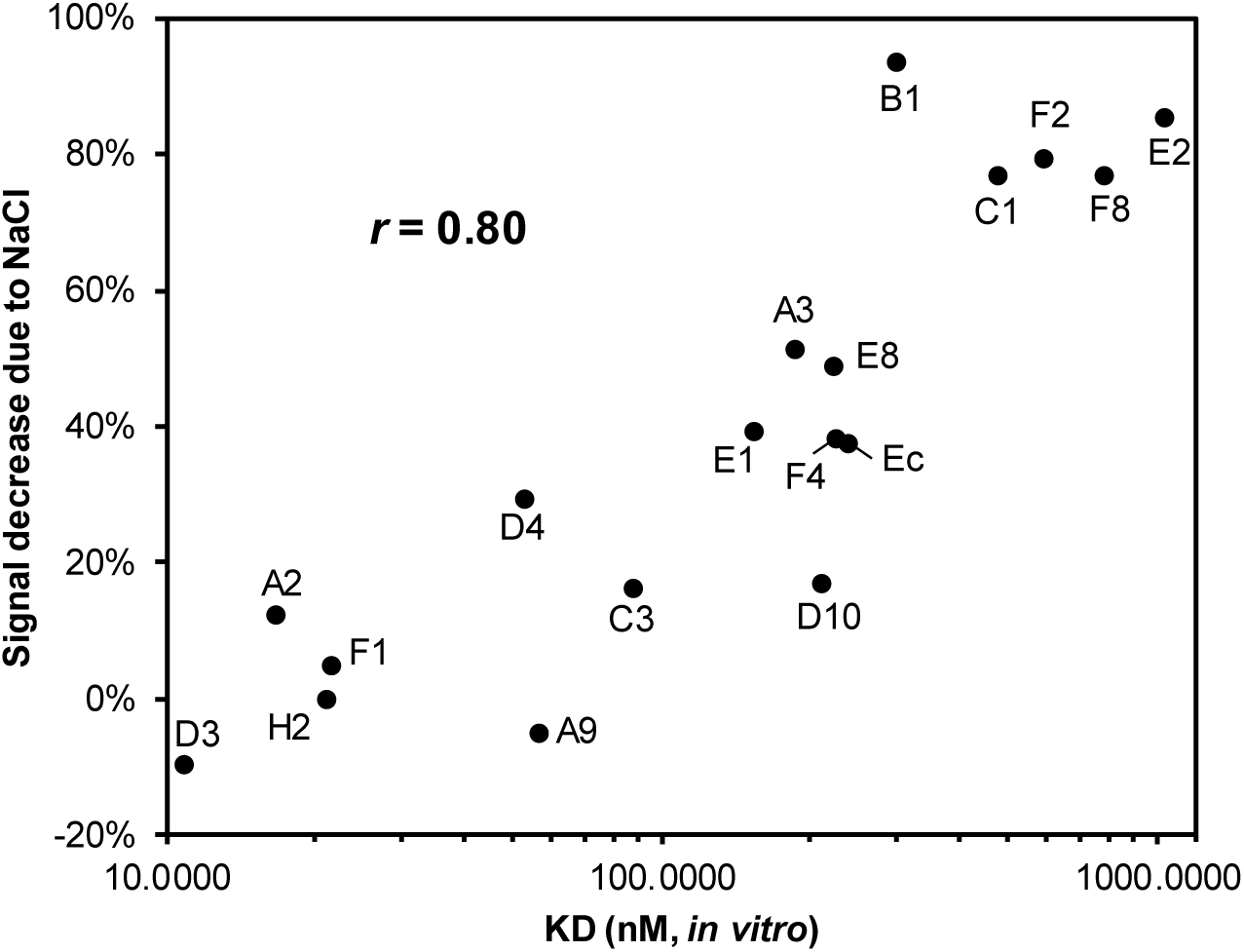
Effect of 67 mM NaCl on the difference between FRET in *ΔpdeH* and *ΔdgcE* backgrounds. The decrease in the FRET difference was more pounced for biosensors with high *K*_D_ values. n=3. Average values are presented. Pearson correlation coefficient *r* is shown.

**Supplementary Fig. 5.**
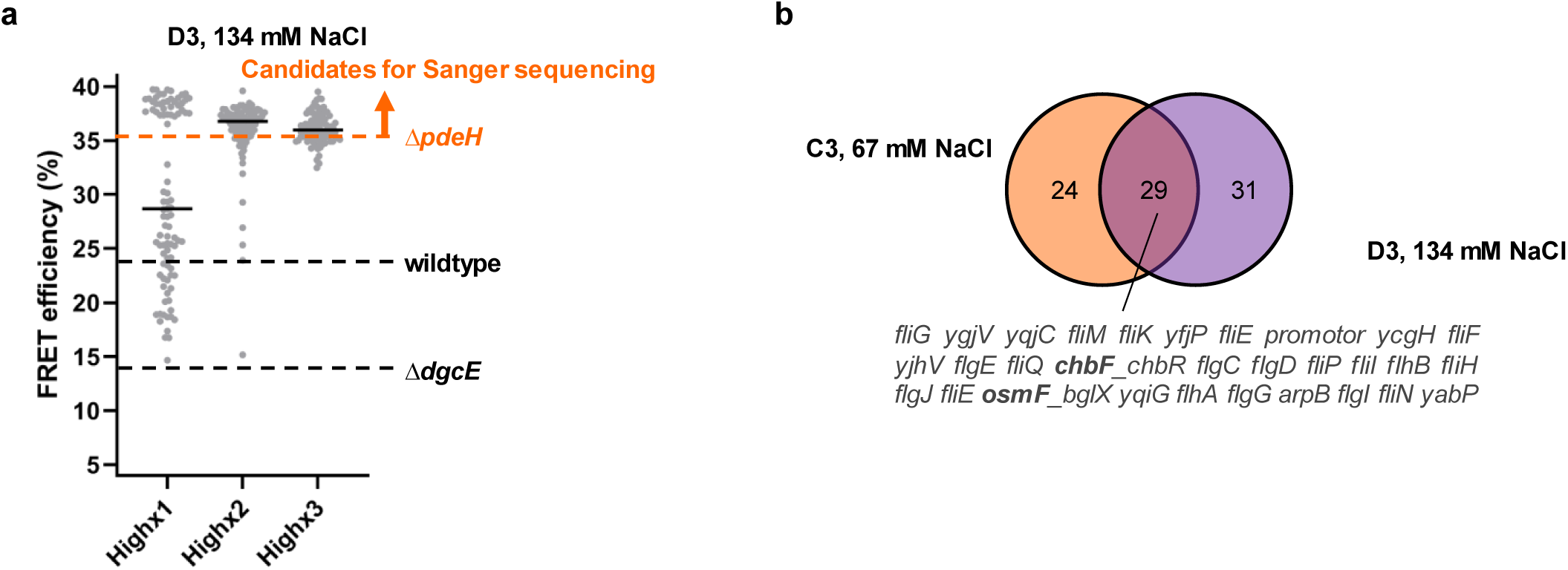
Sorting for mutants with elevated c-di-GMP levels using D3 biosensor in the presence of 134mM NaCl. **a,** Distributions of FRET values for 93 colonies of mutants after the first (x1), second (x2) and third (x3) sorting, respectively, measured using flow cytometry. The D3 biosensor (*K*_D_ ∼11 nM) was utilized to sort mutants with low c-di-GMP levels. The sorting buffer contained 134 mM NaCl, to lower c-di-GMP levels and thus to increase selectivity for high-c-di-GMP sorting. Mutants confirmed to have higher c-di-GMP levels than *ΔpdeH* were genotyped by Sanger sequencing. **b,** Overlap of Sanger sequencing results between the data for high c-di-GMP sorting using C3 biosensor in presence of 67 mM NaCl and D3 biosensor in presence of 134 mM NaCl. Common gene mutations are listed by names. The underline indicates that the insertion located between two neighbor genes. The gene labeled with bold font refers to the insertion is located its upstream, without published information on the promoter region.

**Supplementary Fig. 6.**
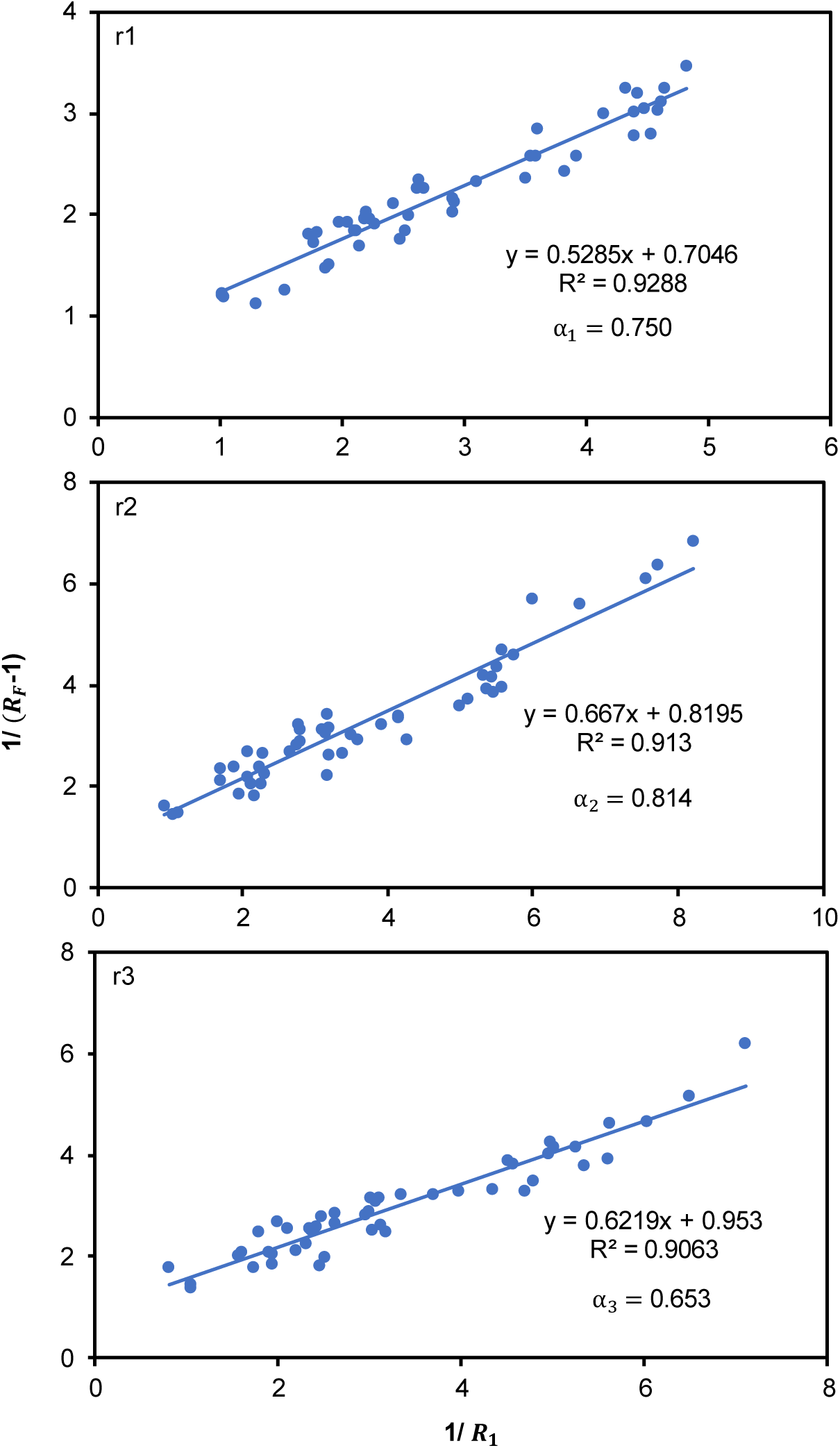
Determination of the calibration factor *α* for the calculation of FRET efficiency from sensitized FRET data obtained using flow cytometry. The linear regressions provide the slope and the intercept from the plot of 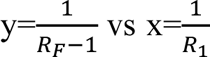. The ratio of the slope and the intercept yields the value of *α*. The r1, r2 and r3 represent independent biological replicates.

**Supplementary Table 1.**
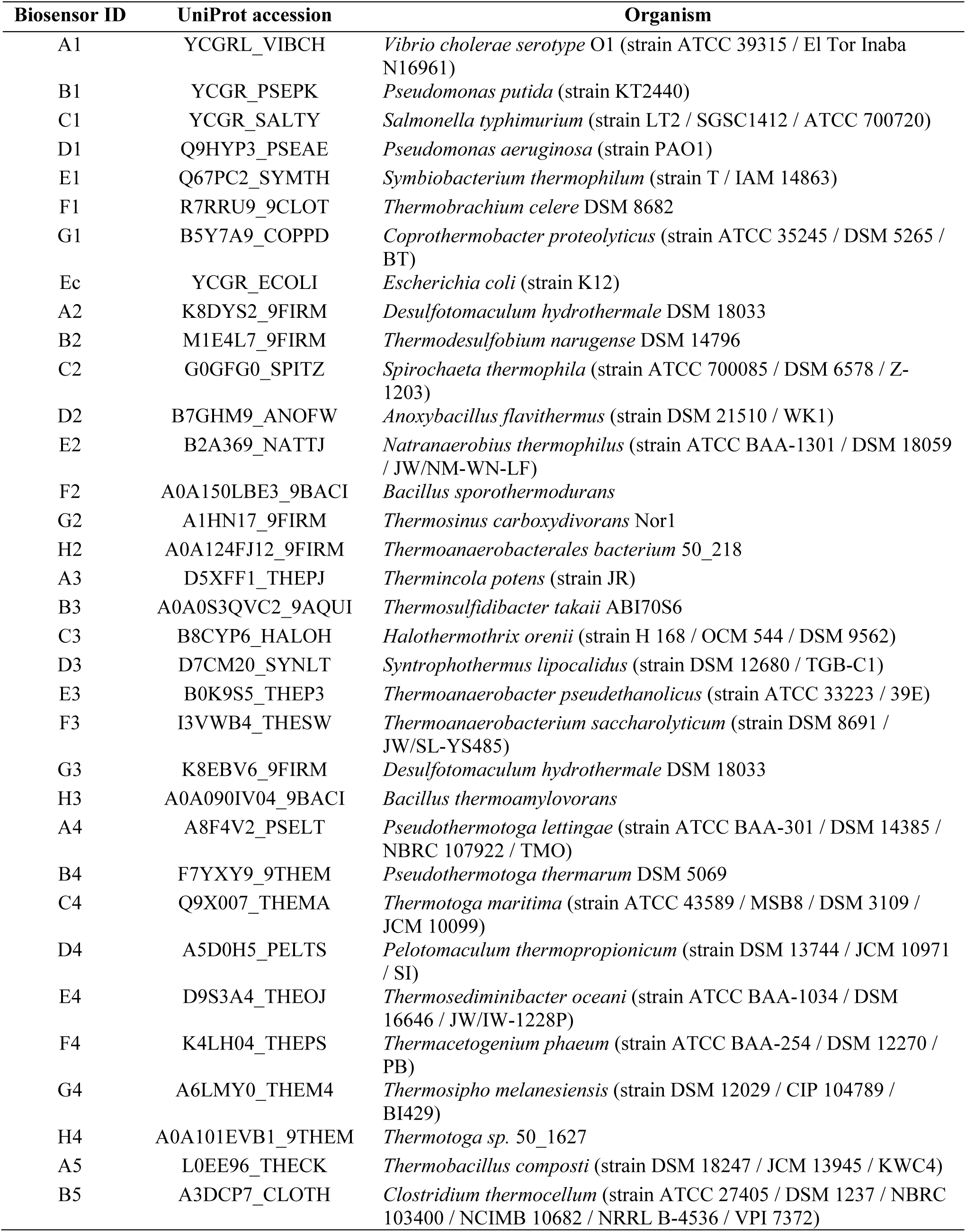
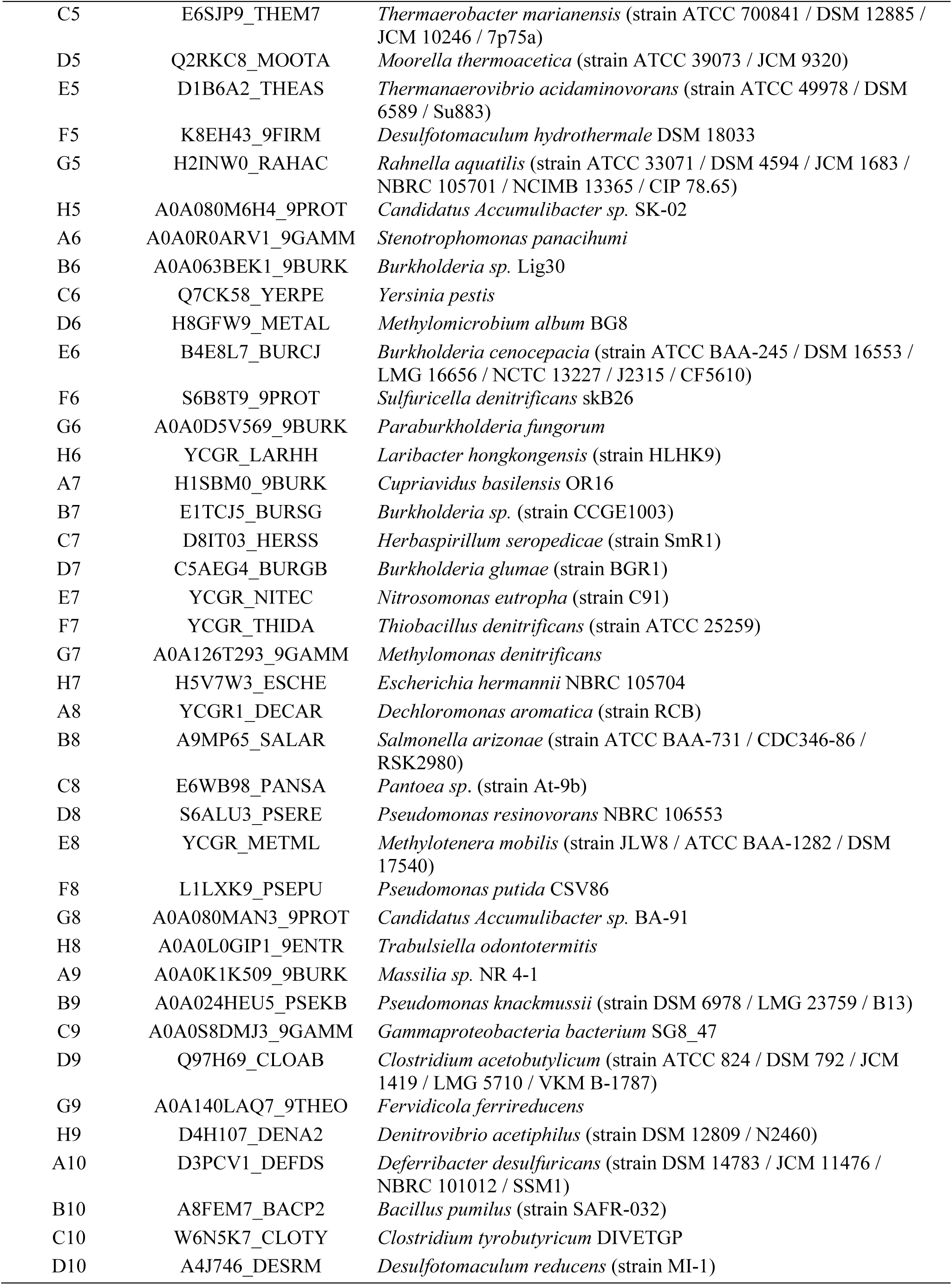
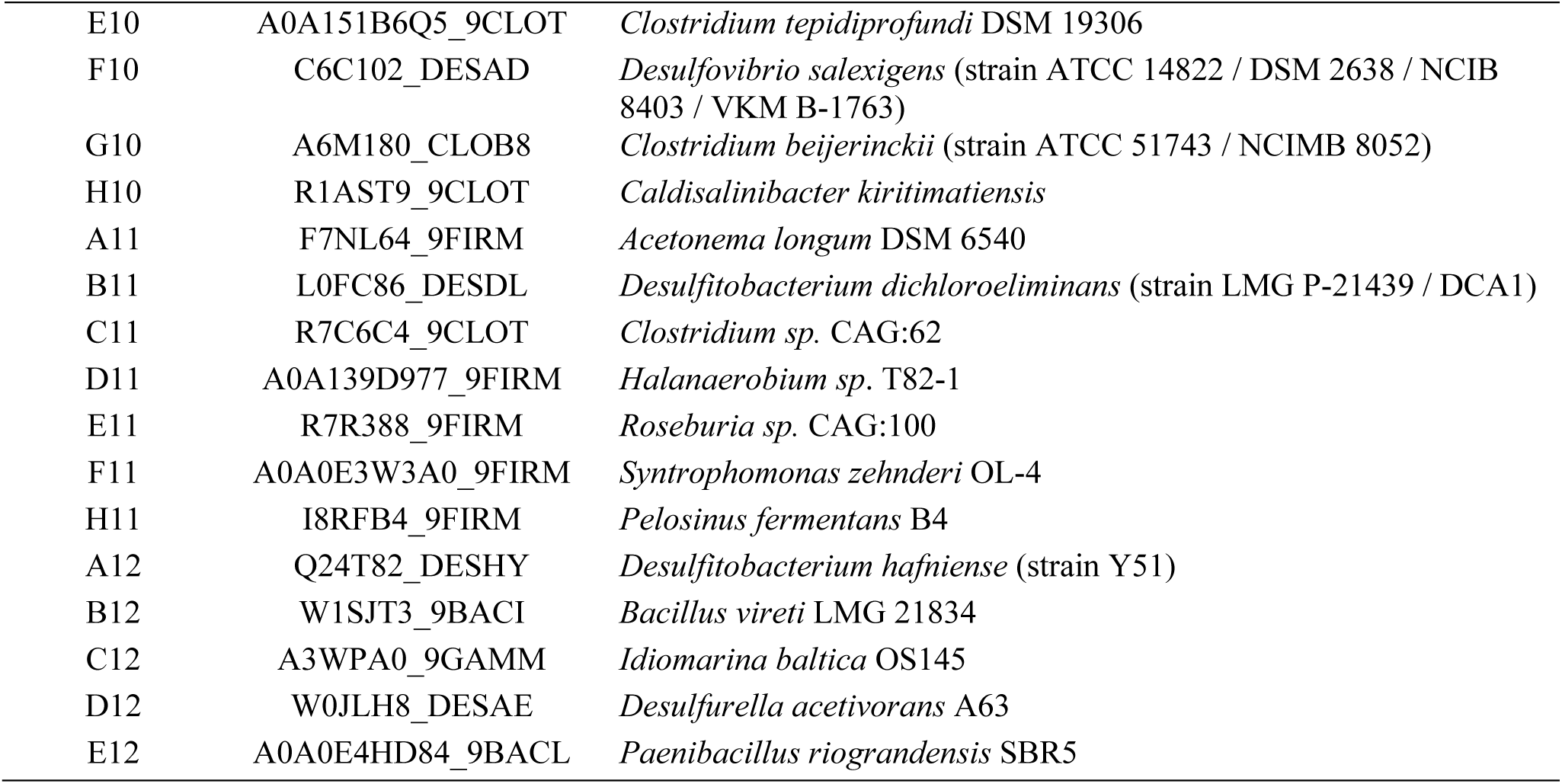
List of PilZ-domain containing proteins used in the biosensor library.

**Supplementary Table 2.**
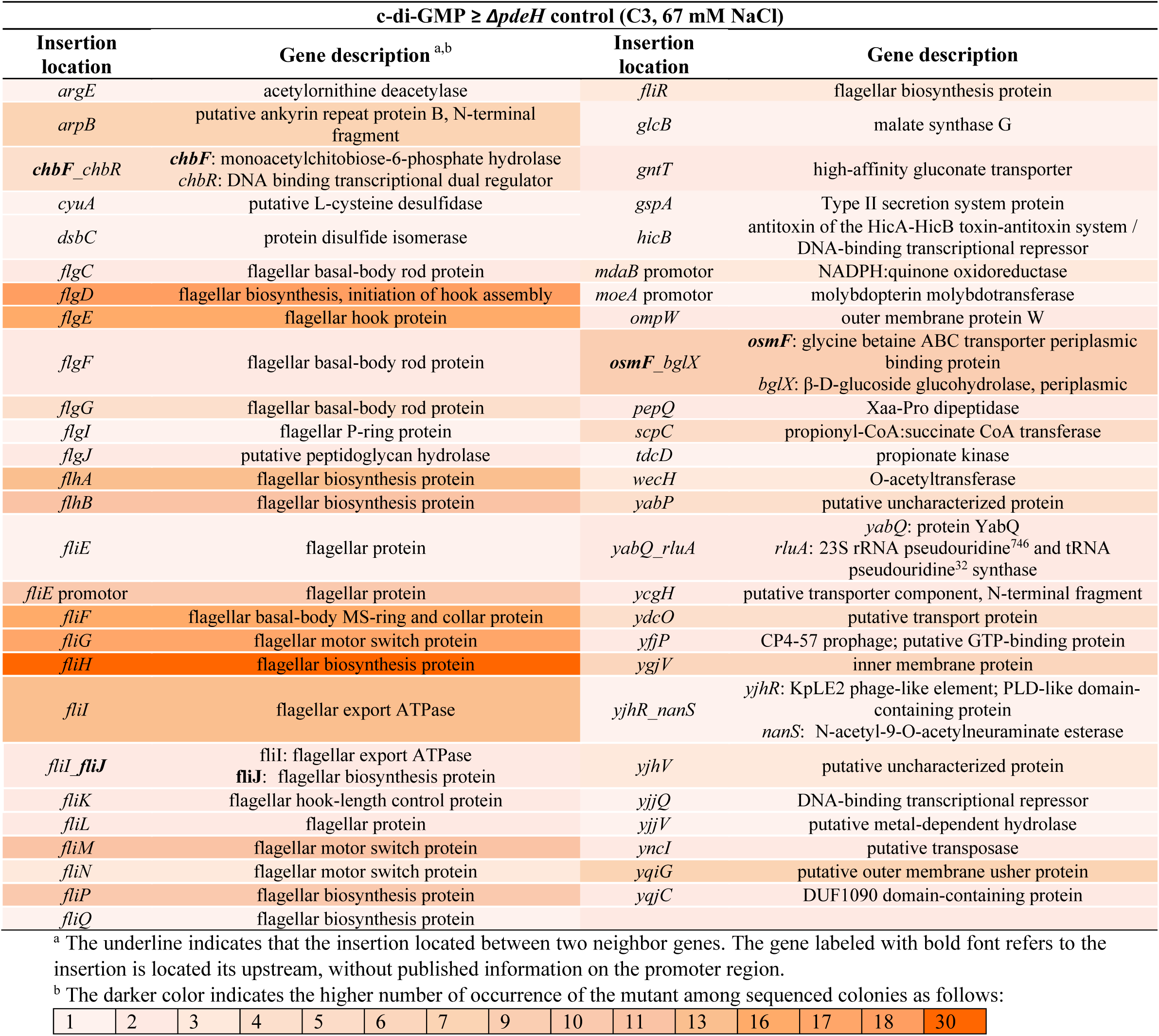
Mutations elevating c-di-GMP levels identified by Sanger sequencing using C3 biosensor.

**Supplementary Table 3.**
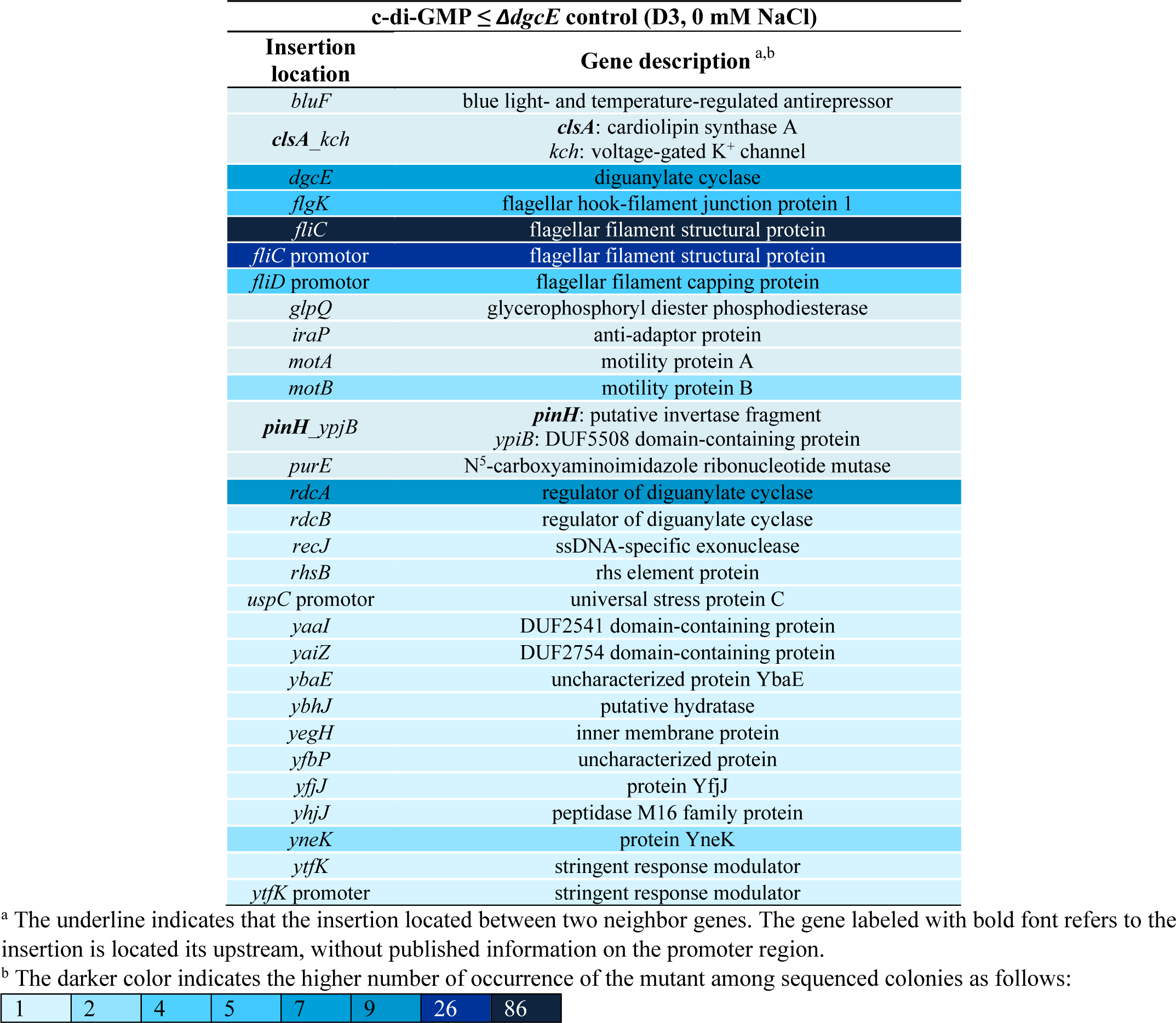
Mutations lowering c-di-GMP levels identified by Sanger sequencing using D3 biosensor.

| **c-di-GMP ≤ ΔdgcE control (D3, 0 mM NaCl)** |  |
| --- | --- |
| **Insertion location** | **Gene description<sup>a,b</sup>** |
| *bluF* | blue light- and temperature-regulated antirepressor |
| *clsA_kch* | *clsA*: cardiolipin synthase A<br>*kch*: voltage-gated K<sup>+</sup> channel |
| *dgcE* | diguanylate cyclase |
| *flgK* | flagellar hook-filament junction protein 1 |
| *fliC* | flagellar filament structural protein |
| *fliC* promotor | flagellar filament structural protein |
| *fliD* promotor | flagellar filament capping protein |
| *glpQ* | glycerophosphoryl diester phosphodiesterase |
| *iraP* | anti-adaptor protein |
| *motA* | motility protein A |
| *motB* | motility protein B |
| *pinH_ypjB* | *pinH*: putative invertase fragment<br>*ypjB*: DUF5508 domain-containing protein |
| *purE* | N<sup>5</sup>-carboxyaminoimidazole ribonucleotide mutase |
| *rdcA* | regulator of diguanylate cyclase |
| *rdcB* | regulator of diguanylate cyclase |
| *recJ* | ssDNA-specific exonuclease |
| *rhsB* | rhs element protein |
| *uspC* promotor | universal stress protein C |
| *yaaI* | DUF2541 domain-containing protein |
| *yaiZ* | DUF2754 domain-containing protein |
| *ybaE* | uncharacterized protein YbaE |
| *ybhJ* | putative hydratase |
| *yegH* | inner membrane protein |
| *yfbP* | uncharacterized protein |
| *yfjJ* | protein YfjJ |
| *yhjJ* | peptidase M16 family protein |
| *yneK* | protein YneK |
| *ytfK* | stringent response modulator |
| *ytfK* promotor | stringent response modulator |

<sup>a</sup> The underline indicates that the insertion located between two neighbor genes. The gene labeled with bold font refers to the insertion is located its upstream, without published information on the promotor region.
<sup>b</sup> The darker color indicates the higher number of occurrence of the mutant among sequenced colonies as follows:

|  |  |  |  |  |  |  |  |
| --- | --- | --- | --- | --- | --- | --- | --- |
| 1 | 2 | 4 | 5 | 7 | 9 | 26 | 86 |

**Supplementary Table 4.**
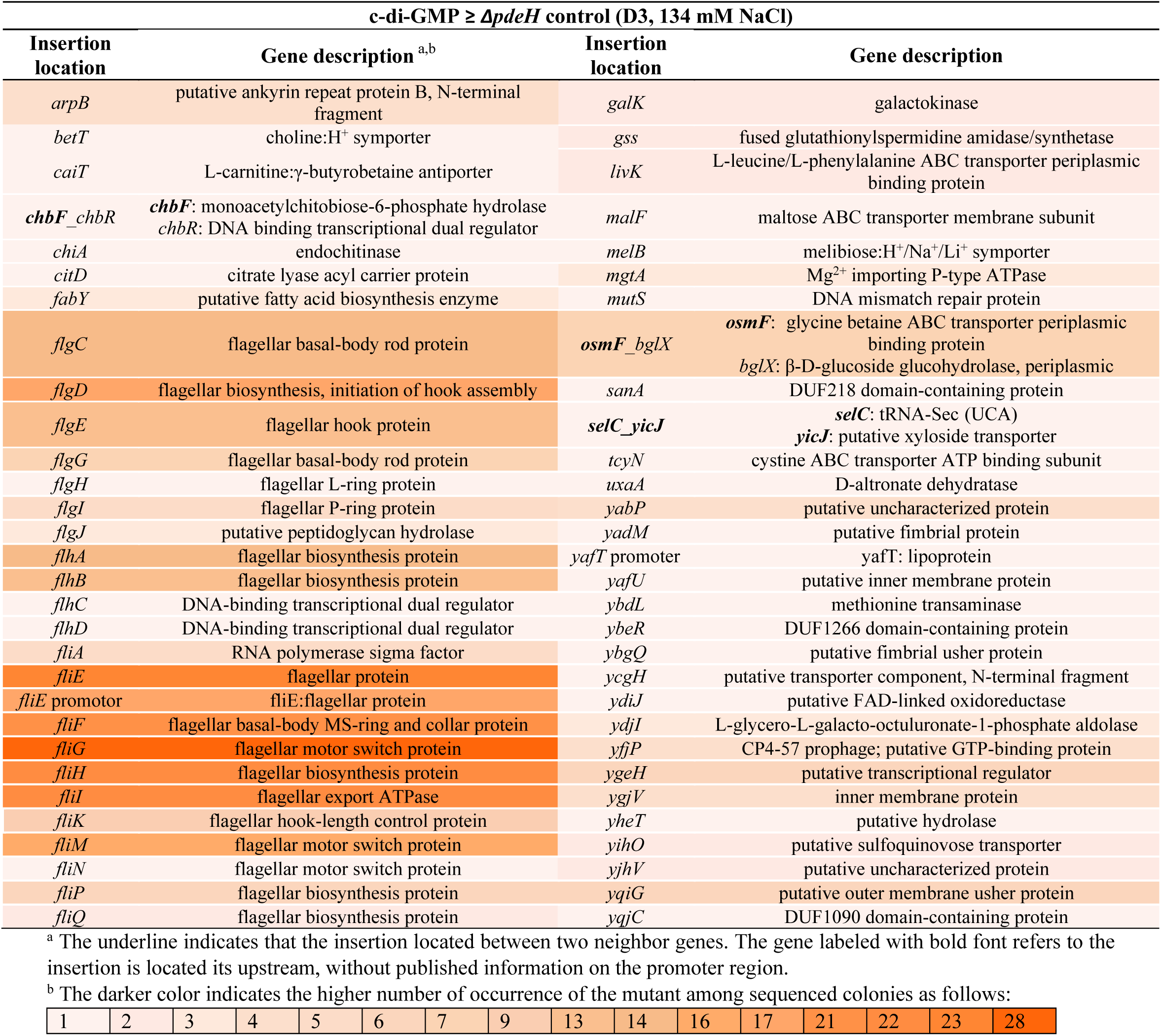
Mutations elevating c-di-GMP levels identified by Sanger sequencing using D3 biosensor.

**Supplementary Table 5.**
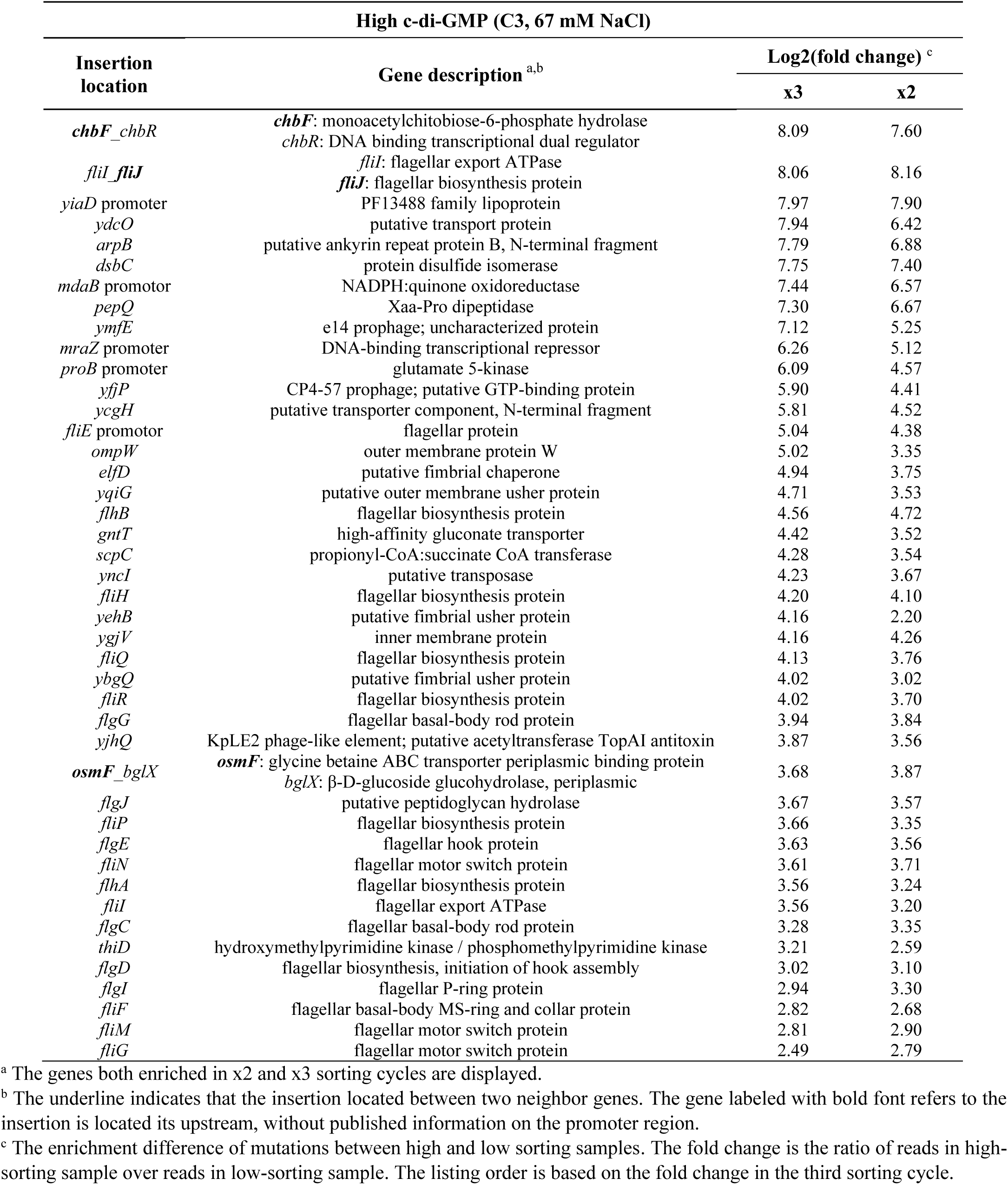
Mutations elevating c-di-GMP levels identified by NGS data using C3 biosensor.

**Supplementary Table 6.**
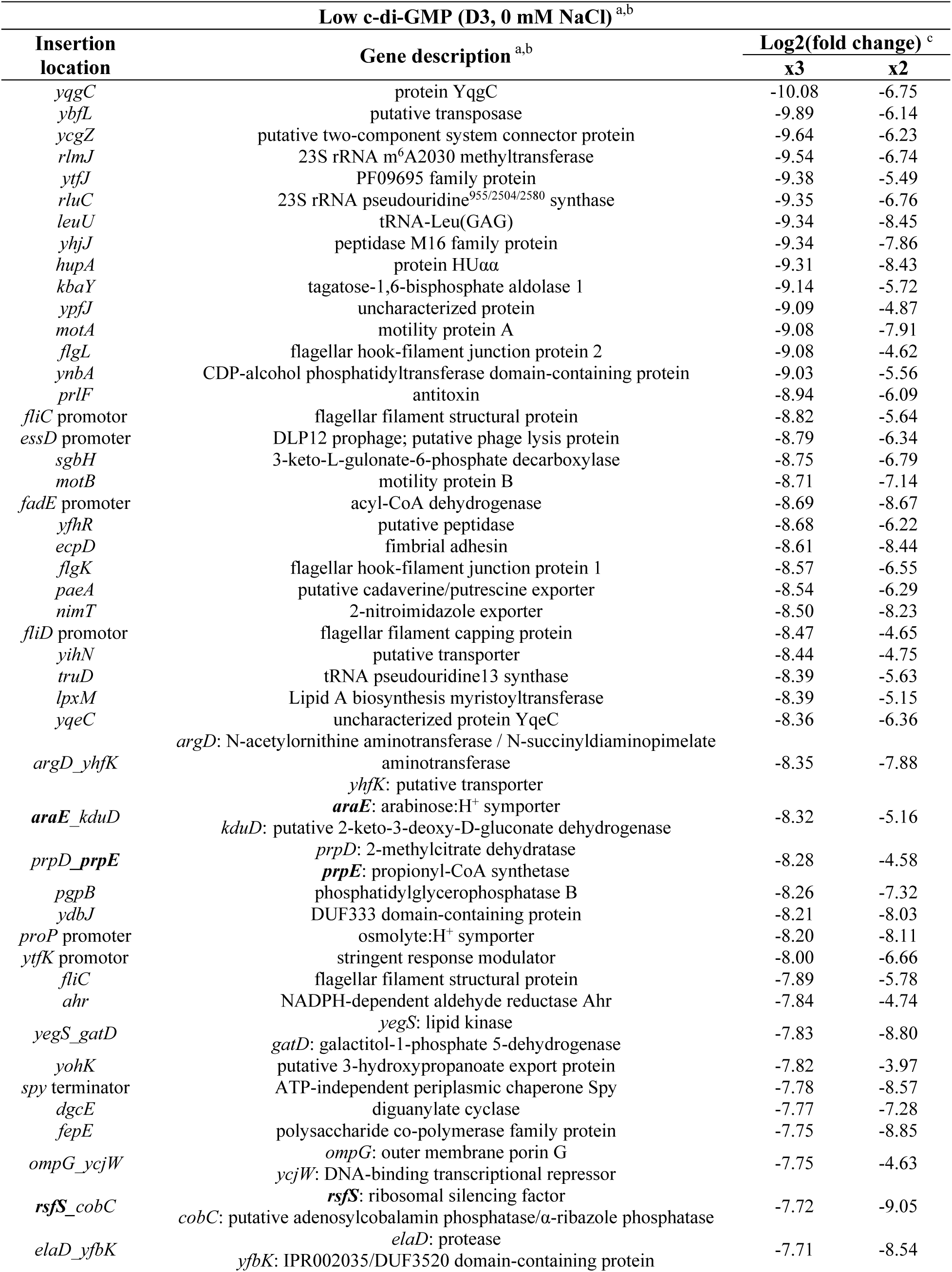
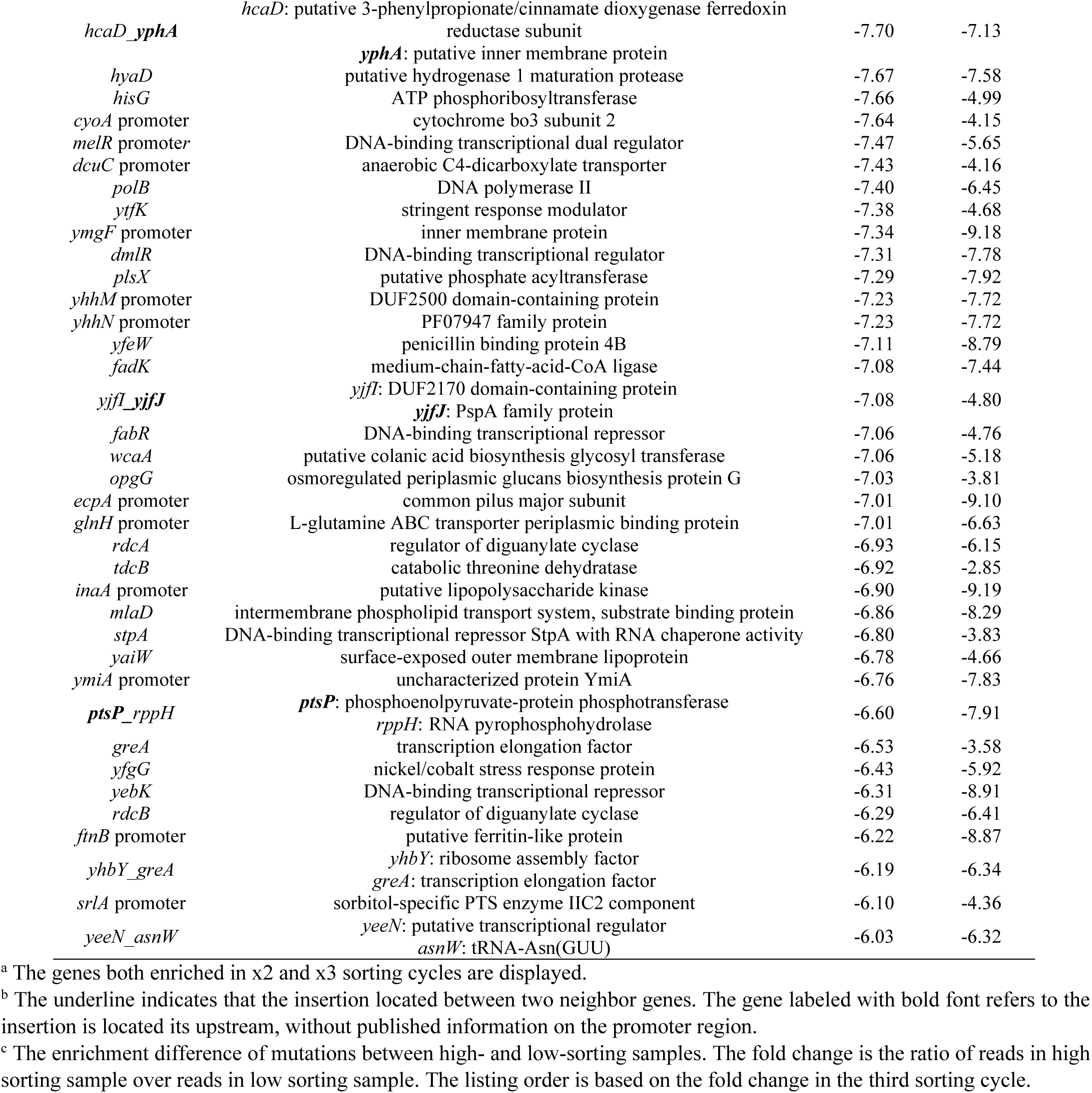
Mutations lowering c-di-GMP levels identified by NGS data using D3 biosensor.

**Supplementary Table 7.** Strains and the plasmids used in this study.

| Strains | Relevant genotype<sup>a</sup> | Reference |
| --- | --- | --- |
| W3110 | Wild-type, derivative with functional RpoS | 1 |
| VS701 | W3110 $\Delta pdeH$ | 2 |
| VS679 | W3110 $\Delta dgcE$ | 2 |
| VS741 | W3110 $\Delta pdeH\Delta dgcE$ | 3 |
| LYW7 | W3110 $\Delta motA$ | This work |
| LYW15 | W3110 $\Delta fliC$ | This work |
| LYW65 | W3110 $\Delta flgM$ | This work |
| LYW66 | W3110 $\Delta flgM\Delta motA$ | This work |
| LYW68 | W3110 $\Delta flgM\Delta fliC$ | This work |
| LYW79 | W3110 $\Delta fliG$ | This work |
| LYW81 | W3110 $\Delta fliH$ | This work |
| LYW85 | W3110 $\Delta flgM\Delta fliG$ | This work |
| LYW87 | W3110 $\Delta flgM\Delta fliH$ | This work |
| LYW102 | W3110 $\Delta pdeH\Delta motA$ | This work |
| LYW104 | W3110 $\Delta pdeH\Delta fliC$ | This work |
| LYW106 | W3110 $\Delta pdeH\Delta dgcE\Delta motA$ | This work |
| LYW108 | W3110 $\Delta pdeH\Delta dgcE\Delta fliC$ | This work |
| **Plasmids** |  |  |
| pTrc99a | Expression vector; IPTG inducible; Amp<sup>R</sup> | 4 |
| pBAD33 | Expression vector; arabinose inducible; Cm<sup>R</sup> | 5 |
| pSIJ8 | Provide arabinose inducible lambda Red recombineering genes and a rhamnose inducible flippase recombinase required for Km<sup>R</sup> removal; Amp<sup>R</sup> | 6 |

<sup>a</sup> Amp<sup>R</sup>, Cm<sup>R</sup> and Km<sup>R</sup> indicate ampicillin, chloramphenicol and kanamycin resistance, respectively.

